# Haplotype-resolved Genome Assemblies Reveal Subgenome Origins, Genome Bias and Reticulate Evolution in Wheatgrass Species

**DOI:** 10.64898/2026.03.27.714782

**Authors:** Yuanyuan Ji, Raju Chaudhary, Nadeem Khan, Sampath Perumal, Zhengping Wang, Kevin C. Koh, Leila Moghaddam Moghanloo, Pierre Hucl, Bill Biligetu, Andrew G. Sharpe, Lingling Jin

## Abstract

Concerns over climate change have intensified the demand for stress resistant crops like hybrid wheatgrass (HWG; *Elymus hoffmannii*, StStStStHH), a perennial forage species known for its exceptional salt and drought tolerance. However, hexaploidy and high heterozygosity have complicated efforts to resolve its genomic structure and evolutionary history. Here, we present high-quality, haplotype-resolved, chromosome-level genome assemblies for HWG (CDC Saltking) and its putative progenitor, bluebunch wheatgrass (*Pseudoroegneria spicata*, StSt). By integrating PacBio HiFi and ultra-long Oxford Nanopore sequencing with Hi-C scaffolding, we assembled the 10.7 Gb HWG genome into 21 pseudochromosomes per haplotype. Our phylogenomic analysis redefines the origin of the H subgenome, positioning it as an intermediate between Old-World *Hordeum marinum* (sea barley) and *Hordeum brevisubulatum*. Notably, we identified significant chromosomal rearrangements, including a unique duplication on St chromosome 4. Transcriptome analysis across multiple tissues revealed a pronounced expression dominance of the H subgenome. This dominance was not associated with reduced LTR density, suggesting that selective pressures for rapid adaptation of the latest subgenome entrant may drive its dominance. Finally, using the *f*-branch statistic, population genomic analysis of 189 accessions representing eight *Elymus* and *Pseudoroegneria* species revealed extensive reticulate evolutionary relationships and identified *P. spicata* as a major, asymmetric genetic donor within the wheatgrass complex. These resources provide a foundational framework for future genomic research and genetic improvement in grasses and for the introgression of stress-tolerance traits into cereal crops such as wheat.

**Key Messages:**

- Development of world-first high-quality chromosomal-level haplotype-resolved genome assemblies of hexaploid HWG and diploid progenitor, *Pseudoroegneria spicata*, enabled the identification of the subgenome origins.
- This study resolved the evolutionary placement of the St genome and clarified the history of polyploidization and hybridization in HWG.
- Homeolog expression bias in the H subgenome likely reflects selective pressure favoring greater gene retention and upregulation of functionally important genes, thereby enhancing hybrid fitness.
- Population structure analysis distinctly differentiates *P. spicata*, *E. repens*, *E. hoffmannii* from other European *Pseudoroegneria* species.
- The findings reveal the complex patterns of interspecific gene flow and population dynamics within the *Elymus* and *Pseudoroegneria* species.

## Introduction

Escalating climate change have driven an increased demand for resilient crop varieties with high adaptability to barren soils and abiotic stress conditions such as drought and salinity^1–3^. Hybrid wheatgrass (*Elymus hoffmannii*, StStStStHH, HWG), is a perennial forage grass recognized for its strong tolerance to salinity and drought, offering a sustainable solution for livestock production while reducing soil erosion^4–6^. Although various breeding strategies have been proposed to improve the yield and quality of forage wheatgrasses^7,8^, achieving stable yield under highly variable, resource constrained environments remains a major challenge. In this context, generating a high-quality genome sequence of HWG and deciphering its genomic architecture and evolutionary history have become urgent priorities for making a breakthrough in genetic improvement.

One of the primary challenges in assembling polyploid plant genomes is their inherent genomic complexity, which largely arises from recurrent polyploidization and interspecific hybridization events^9–11^. This challenge is particularly pronounced in HWG, an outcrossing hexaploid grass that exhibits high levels of heterozygosity. To date, four cultivated varieties have been developed in this grass, including AC Saltlander^12^, NewHy^13^, RS Hybrid^14^, and CDC Saltking^5^, all of which share the same StStStStHH genomic constitution^15^.

Polyploid species are generally classified as auto- or allopolyploid based on their origin. Autopolyploids arise through genome duplication within a single diploid species, while allopolyploids result from genome doubling following the hybridization between two distinct diploid progenitors^16^. Due to the absence of high-quality genome sequences for HWG and its diploid or tetraploid relatives, the evolutionary origin of the two St subgenomes remains unresolved. In particular, it is unclear whether HWG represents an autopolyploid or an allopolyploid with respect to its St subgenome composition. Because polyploidization and hybridization are major evolutionary forces underlying the emergence of agronomically important traits and adaptation to diverse environments^17,18^, elucidating the history of these events among the three subgenomes of HWG will offer valuable insights into its genetic potential for forage crop improvement and facilitate the introgression of desirable traits into related crops such as wheat.

In this study, we report two high-quality, haplotype-resolved genome assemblies: one for HWG (*Elymus hoffmannii*, cultivar CDC Salt king) and one for its putative progenitor, *Pseudoroegneria spicata* (PI635993). By comparing these two newly assembled genomes with other publicly available genomic resources, we aim to: (1) resolve the evolutionary position of HWG within the Poaceae family; (2) uncover its polyploidization and hybridization history; (3) identify the closest extant relatives of its diploid progenitors; (4) characterize patterns of subgenome dominance and their genetic basis; and (5) examine population structure and introgression events within *Elymus* and *Pseudoroegneria* species.

## Results

### Chromosome-Level Haplotype-Resolved Assemblies of *E. hoffmannii* and *P. spicata*

We generated haplotype-resolved, chromosome-level genome assemblies for HWG (*E. hoffmannii,* 2n = 6x = 42) and one of its progenitor *P. spicata* (PI635993, 2n = 14)^19^ using an integrated sequencing and scaffolding strategy that combined Pacbio HiFi sequencing, Hi-C sequencing, and Oxford Nanopore ultra-long read sequencing. Initial flow cytometry analyses estimated haploid genome sizes of 11.12 Gb for *E. hoffmannii* and 3.2 Gb for *P. spicata* (Supplementary Figure 2a). The draft HWG genome assembly was partitioned into two haplotypes (hap1 and hap2), with total sizes of 10.7 Gb and 10.5 Gb, respectively. Each haplotype was scaffolded into 21 chromosomes using Hi-C data, achieving scaffold N50 values of 436.71 Mb for hap1 and 463.45 Mb for hap2. Similarly, the *P. spicata* genome was assembled into two haplotypes of 3.77 Gb and 3.72 Gb, with corresponding scaffold N50 values of 534.39 Mb and 502.69 Mb (Table 1). For both species, the haploid genome assembly sizes were consistent with estimates obtained from flow cytometry (Supplementary Figure 2a).

**Table 1:**
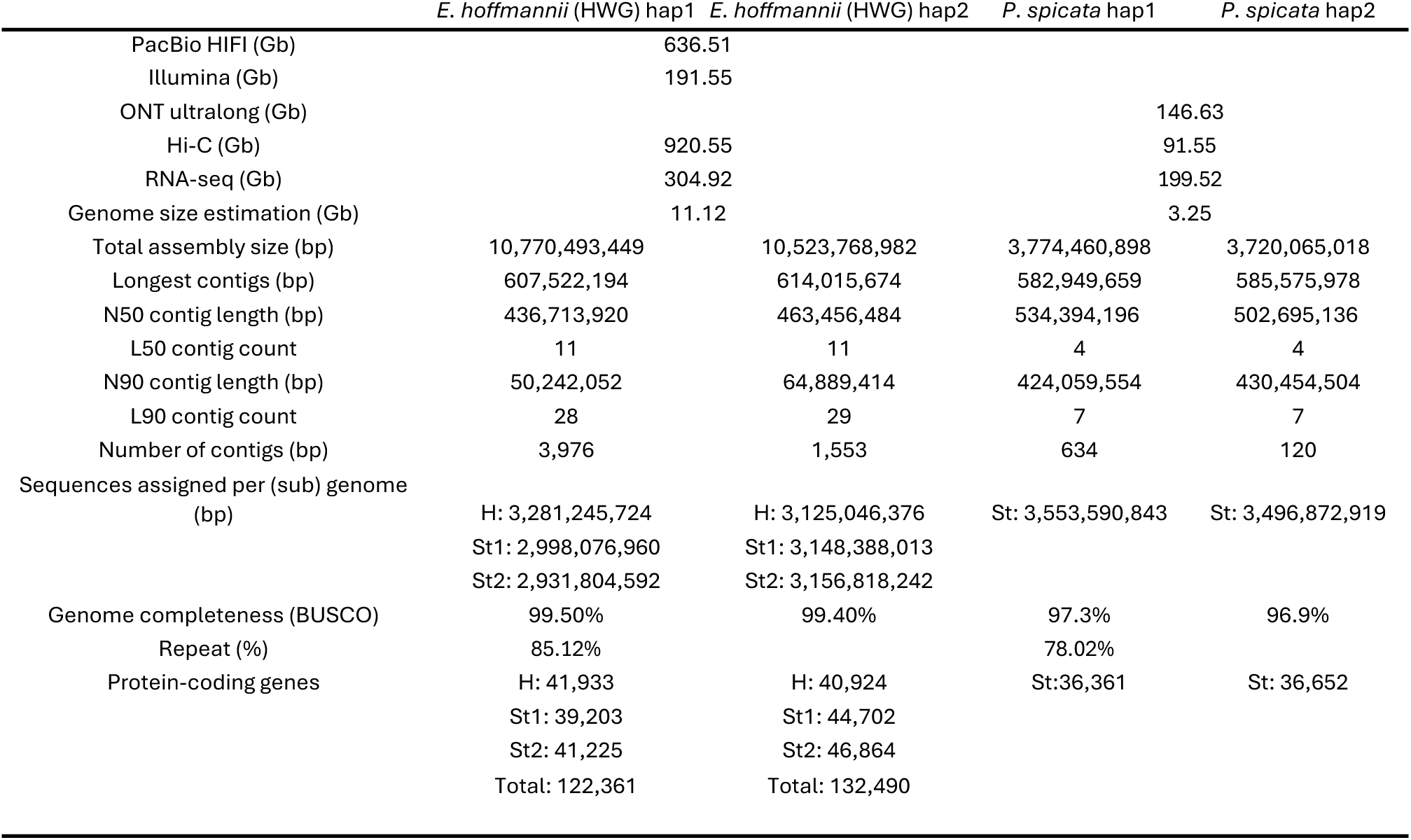
Genome assembly statistics for diploid and hexaploid hybrid wheatgrass species.

Given the complex polyploid genome architecture containing two St subgenomes, we employed an allele-aware scaffolding tool HapHiC^20^ to construct pseudochromosomes. Using this approach, 91.4% and 93.8% of the assembled contigs were anchored to 21 pseudochromosomes for each haplotype, respectively. Hi-C interaction patterns showed strong concordance between homologous chromosomes, indicating a high degree of homology in subgenomes, particularly between the two St genomes (Supplementary Figure 2b). Assembly quality was further supported by benchmarking metrics, including a BUSCO completeness score of 99.0% and long terminal repeat (LTR) assembly indices (LAI) (LAI_H-mean_=19.15, LAI_St1-mean_=17.24, LAI_St2-mean_=18.66) (Table 1 and Supplementary Figure 4c). For *P. spicata*, 94.1% and 94.0% of assembled contigs were anchored to 7 pseudochromosomes in hap 1 and hap 2, respectively. These assemblies also exhibited high completeness, with BUSCO scores exceeding 96.9% and a mean LAI of 19.64, consistent with reference-level scaffolding (Supplementary Figure 4d). Comparative collinearity analysis between barley (*H. vulgare*) and *P. spicata* identified a chromosomal inversion in hap2, whereas hap1 showed no such structural rearrangement (Supplementary Figure 3d).

We annotated all genomes using a combination of *ab initio* and RNA-seq-based gene prediction approaches. In HWG, the final gene counts for hap1 and 2 are 41,933/40,924 in the H subgenome, 39,203/44,702 in St1, and 41,225/46,864 in St2 respectively (Table 1). For *P. spicata*, a total of 36,361 and 36,652 protein coding genes were identified for hap1 and 2, respectively (Table 1). These gene numbers are comparable to those reported for closely related species, including *Thinopyrum elongatum* (44,474)^21^ and tetraploid *Elymus sibiricus* (89,800)^22^. Genome completeness was further supported by BUSCO analysis, which showed that more than 93% of conserved BUSCO genes were complete in both genomes (Table 1 and Supplementary Figure 4b).

### Chromosome Origins and Subgenome Assignment in HWG

Among the four varieties, AC Saltlander is believed to derived from a natural crossing between quackgrass (*Elymus repens*, 2n = 42, StStStStHH) and a *Pseudoroegneria* species carrying the St genome^12^, whereas, Newhy, RS hybrid and CDC Saltking are synthetic hybrids generated through crosses involving quackgrass or AC Saltlander and *P. spicata*^4,5,13,14^. Despite their distinct backgrounds, all four varieties share key phenotypic improvements over their quackgrass ancestor, including enhanced tolerance to salinity and reduced rhizome^12–15^. Because the H genome is directly inherited from *E. repens* without additional introgression, the observed phenotypic differences are more likely driven by divergence within the St genomes. Although previous cytological evidence suggests a high similarity between the two St genomes^15^. The classification of the two St genome sets in HWG as either allopolyploid or autopolyploid remains debated.

To accurately assign chromosomes to their respective subgenomes in the hexaploid HWG, we first used a *k*-mer-based method to identify subgenome-specific sequence signatures^23^. Based on differential *k*-mer profiles, the 21 chromosomes in hap1 were grouped into three clusters comprising 7, 4, and 10 chromosomes (Figure 1b). The uneven chromosome distribution suggests that high sequence homology between the two St subgenomes blurs their *k*-mer signatures, resulting in the misassignment to the incorrect subgenome.

**Fig. 1.**
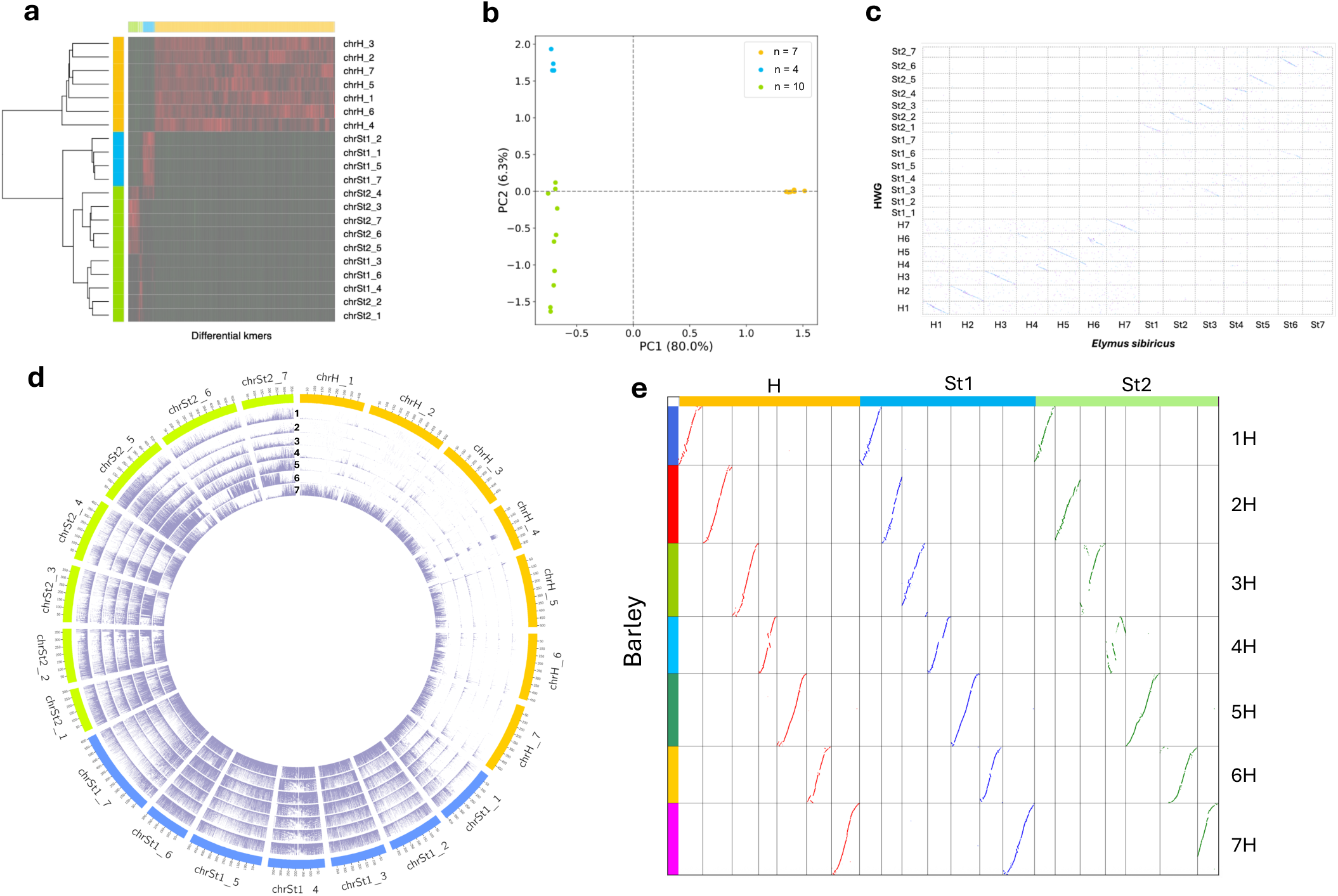
Subgenome chromosome assignment of HWG and tracing the origin of chromosomes. **a,** Clustering of differential *k*-mers (*k* = 15 and frequency ≥ 50) among homoeologous chromosomes. **b**, Principal components analysis of subgenome chromosomes clustered based on differential *k*-mers. The n values for each clustering indicate the number of chromosomes that clustered in the same group. **c**. Dot plot shows the identity of each chromosomes using *E. sibiricus* as a reference genome. Each dot represent a region of DNA alignments between HWG and *E. sibiricus*. A minimum length of 200 bp and cluster size of 100 were used in dotplot analysis. **d**. circos plot shows the coverage of GBS sequencing reads mapping to HWG (hap1) genome. Track 1: *P. libanotica* (PI228389), Track 2: *P. tauri* (PI380652), Track 3: *P. strigosa* (PI499637), Track 4: *P. geniculata* (PI692218), Track 5: *P. stipifolia* (PI675310), Track 6: *P. spicata* (PI635993), Track 7: *E. repens* (PI565007). **e**, Collinearity analysis comparing HWG (hap1) homologous chromosomes and Barley (*H. vuglare*) chromosomes.

Alternatively, dotplot comparisons between HWG and its tetraploid relative *E. sibiricus* (2n=4x=28, StStHH)^22^ identified the seven H chromosomes and three misclassified St chromosomes (Chr3, Chr4, Chr6) (Figure 1c). Validation against the diploid progenitor *P. spicata* using sensitive alignment parameters confirmed that these three chromosomes belong to the St_1_ subgenome (Supplementary Figure 8). Furthermore, collinearity analysis with barley validated the overall chromosome assignments (Figure 1e). Notably, Chr4 of the St_2_ subgenome displayed an overlapping collinear region, indicating a chromosomal duplication that consistently observed across both HWG haplotypes (Supplementary Figure 9).

To confirm the chromosome origins of HWG and identify its parental genomes, we performed genotype-by-sequencing (GBS) on seven species including six *Pseudoroegneria* species (StSt) and one *E. repens* (StStStStHH) and mapped GBS reads to the HWG hap1 genome. Among the *Pseudoroegneria* species, only *P. spicata* (StSt) exhibited a complementary mapping pattern with *E. repens*, providing strong evidence that HWG originated from hybridization between these two species (Figure 1d).

### Genomic Structures and Transposable Element (TE) Dynamics in Highly Heterozygous HWG Genome

Chromosomal rearrangements have been suggested to play a major role in plant adaptation and speciation^24,25^. Consistent with this, our comparative structural variation analysis of the HWG, barley and *P. spicata* genomes revealed extensive large-scale rearrangements through HWG evolution. We identified fourteen large inversions (> 50 Mb) and 4 major translocations (> 15 Mb), specifically involving Chr1 and Chr3 of St_1_ and Chr3, Chr4, Chr6 of St_2_ (Figure 2b). These rearrangements collectively affect 3,943 genes (H:658, St_1_:437, St_2_:2,848).

**Fig. 2.**
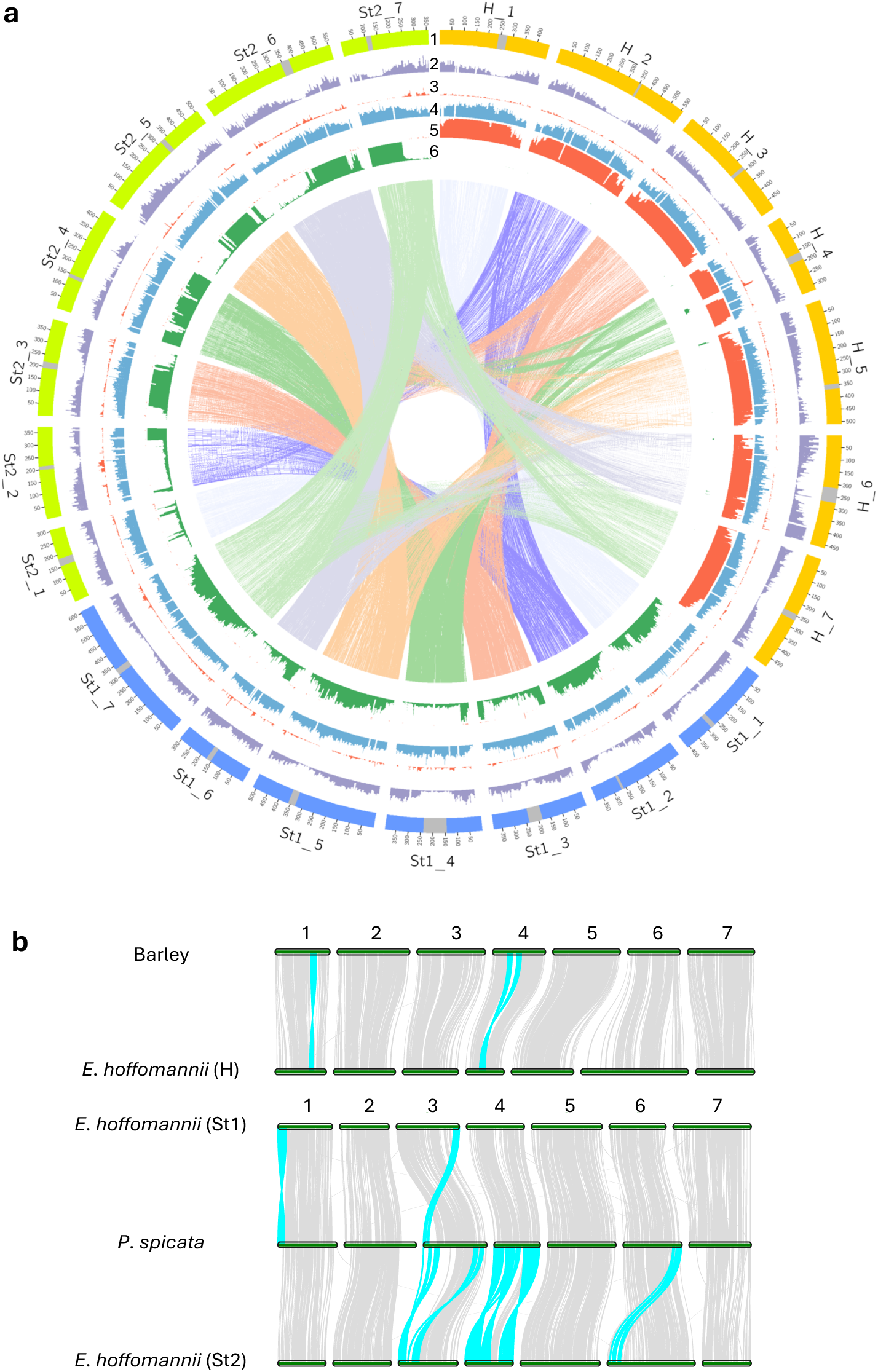
Overview of genomic features (a) and subgenome structure (b) of the assembled hexaploidy *Elymus hoffmannii* genome (hap1). **a.** (1). Centromere position of the H, St_1_, and St_2_ genome chromosomes. (2). The density of protein-coding genes along each chromosome. (3). Non-reference genomic variants distributions. (4). LTR retrotransposon density. (5). H subgenome specific *k*-mer frequencies. (6). St subgenome specific *k*-mer frequencies. The innermost layer shows synteny between subgenome chromosomes. **b**. Structural variations between H genomes and St genomes, large translocations and inversions were highlighted in turquoise.

Across its three subgenomes, HWG shows a total TE content of 85.12%, slightly higher than that of *E. sibiricus* (82.49%)^22^ and significantly higher than its diploid ancestor, *P. spicata* (78.02%) (Supplementary Table S1 and S2). LTR elements were the most abundant repeat type, accounting for 62.88% to 67.21% of the three subgenomes, with the highest proportion observed in St_1_. Among DNA transposons, CACTA was the most enriched superfamily across all three sbugenomes. In contrast, long interspersed nuclear element (LINE) content varied among subgenomes, comprising 1.44% in H, 1.13% in St1, and 2.66% in St2. Moreover, TE composition differs markedly between the St genomes of *P. spicata* and HWG. In *P. spicata*, TIR elements occur at substantially lower copy numbers than in HWG St genomes, yet exhibit greater diversity in TIR types (Supplementary Table 1). An overview of genome-wide TE distribution in HWG is provided in Supplementary Figure 6.

In addition, to evaluate the heterozygosity of HWG, we mapped Illumina paired-end sequencing reads to the HWG hap1 assembly, identifying 48,110,935 high-quality variants. The resulting heterozygosity rate of 0.45% (heterozygous SNPs and InDels: 46,164,944 and 1,388,819) is significantly higher than that of related grasses, such as wheat ( < 0.1%)^26,27^ and oat (0.011%)^10^. To characterize the genomic variations between hap1 and hap2, we identified 16,817 (0.036%) and 4,673 (0.336%) non-reference heterozygous SNPs and InDels, using hap1 as the exclusive reference genome for mapping. These non-reference heterozygous SNPs and InDels represent the specific points where hap2 deviates from hap1, highlighting substantial sequence divergence between hap1 and hap2 (Figure 2a: Track 3).

### Evolutionary Placement of HWG Subgenomes Among Cereal Crops and Its Polyploidization History

Similar to the H subgenome in barley, the St subgenome in *Pseudoroegneria* species also proposed to be evolutionarily ancient^28^. This genomic composition makes HWG particularly informative for subgenome evolution within Triticeae. To clarify the evolutionary positions of the HWG and *P. spicata* subgenomes and their relationship to other sequenced Triticeae genomes, we conducted a gene family analysis across 16 subgenomes in 11 species from the Poaceae family identifying 975 single-copy orthologs. Phylogenetic relationships inferred from sequence alignments of these orthologs using a maximum likelihood method (Figure 3a) showed that the H subgenome in HWG and *E. sibiricus* diverged approximately 8.2 Mya and are more closely related to *H. marinum* than to *H. vulgare* within the Hordeum clade. The St subgenomes diversified around 17.6 Mya—slightly later than Hordeum (19.75 Mya) but earlier than the *Secale cereale* (15.18 Mya) and *Thinopyrum elongatum* (12.66 Mya). The divergence between St1 and St2 genomes was estimated at approximately 14.23 Mya, with *P. spicata* showing closer genetic affinity to St2. These divergence times and genomic relationships between the H and St lineages were further supported by synonymous substitutions (*K*s) rate analyses of orthologous coding genes (Figure 3c).

**Fig. 3.**
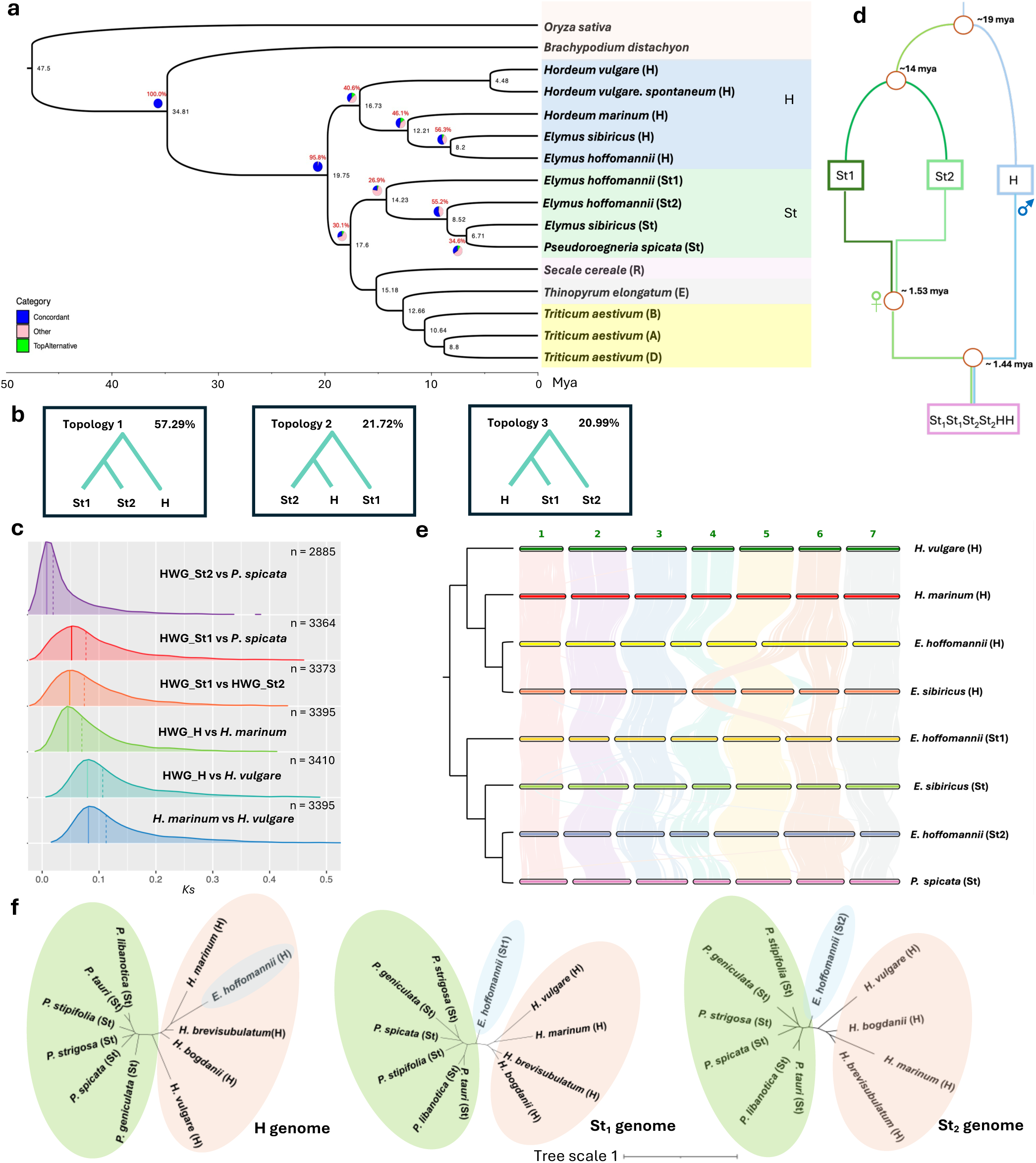
Phylogenetic relationship of 16 genomes, including H, St_1_, St_2_ and public sequenced genomes in Poaceae. **a**, The phylogenetic tree was reconstructed from 975 single-copy genes. The divergence times among different genomes are labelled on the branch point. Pie charts along the backbone phylogeny demonstrate the proportion of gene trees supporting the represented topology. **b**, Proportions of conflicting gene tree topologies among the three major topological patterns observed for the H, St_1_, and St_2_ subgenomes, based on 975 single-copy genes. **c**. Divergence time was estimated based on *K*s distributions. Density distributions of *K*s values are derived from pairwise comparisons of syntologs between different genomes of each species. Solid line indicates the peak of the curve, Dash line represent the median value of the density distribution. **d**. Model showing the evolutionary history of hexaploid HWG (St_1_St_1_St_2_St_2_HH). Divergence time is determined based on phylogenetic tree in Fig 3a, hybridization time is roughly estimated based on LTR insertion time. **e**. Chromosome evolution and synteny between the subgenomes of HWG, *E. sibiricus*, *P. spicata*, *H. marinum*, and *H. vulgare*. **f**, Unrooted phylogenetic tree of the St and H lineages constructed using

To further identify the donors of H and St subgenomes, we analyzed a combination of publicly available whole-genome sequencing (WGS) data and our GBS datasets, including 10 lineages that represent diverse genomic subtypes (Methods). In total, 286.45 Gb of sequencing data were mapped to the H, St_1_ and St_2_ subgenomes, and unrooted phylogenetic trees were reconstructed using SNPs variants. These analyses showed that the H subgenome is more closely related to *H. marinum* and *H. brevisubulatum* than to *H. vulgare*, whereas both St_1_ and St_2_ subgenomes occupy an intermediate position between *H. vulgare* and *Pseudoroegneria* species (Figure 3f). To further explore subgenome-specific evolutionary patterns, we constructed phylogenetic trees using single-copy genes grouped by chromosomes. Most chromosomes supported the genome-wide consensus topology; however, Chr4 of the St_2_ subgenome displayed an earlier divergence relative to its St_1_ counterpart (Supplementary Figure 10), making it the only chromosome that deviated from the overall pattern. In addition, we examined syntenic relationships among *Elymus* species, including HWG, *E. sibiricus*, and their diploid progenitors (*H. vulgare*, *H. marinum* and *P. spicata*). Overall, synteny was largely conserved among the St subgenomes, with only a few large-scale structural rearrangements in Chr 3, 4, and 6. In contrast, a large region of translocation between Chr4 and Chr6 was detected in the H subgenome of *E. sibiricus* but was absent in HWG, indicating an independent evolutionary history of HWG (Figure 3e).

Given the wide distribution of tetraploid wheatgrass such as *E. sibiricus* (StStHH) and *Pseudoroegneria* species (StStStSt), an important question is whether HWG originated from an initial tetraploid of *Pseudoroegneria* species or from an allotetraploid StStHH ancestor. To address this aspect of HWG polyploidization history, we focus on major gene tree conflicts involving the H, St1 and St2 subgenomes. The predominant topology, ((St1,St2), H), accounted for 57.29% of gene trees, whereas the alternative topologies—((St2,H), St1) and ((H,St1), St2)—were observed at much lower and nearly equal frequencies (21.72% and 20.99%, respectively Figure 3b). The prevalence of ((St1,St2), H) topology (>50%) supports a stepwise polyploid origin, where St1 and St2 initially hybridized, followed by a second hybridization with the H genome. Consistent with expectations under the incomplete lineage sorting (ILS) model, the two alternative topologies occur at similar frequencies (∼21%), suggesting a short interval between divergence of the St lineages and their subsequent hybridization with the H genome. The lack of frequency bias between these minority topologies further suggests that the observed phylogenetic discordance is more likely due to the stochastic sorting of ancestral polymorphisms rather than asymmetric gene flow^29^. To independently assess the order of hybridization events, we examined LTR insertion times across subgenomes and found that LTRs in the H subgenome are consistently younger than those in the St subgenomes, providing further support to this evolutionary scenario (Supplementary Figure 12).

Furthermore, the plastid phylogeny was reconstructed using whole-chloroplast genome alignment from 10 species. The results showed clear maternal inheritance of St genome during the second polyploidization event (Supplementary Figure 11). Based on these results, we propose a refined model for the polyploidization history of HWG (Figure 3d), in which two diverged St genomes first hybridized to form an allotetraploid (St1St1St2St2). This tetraploid species subsequently acted as the maternal donor in a second hybridization with H genomes, giving rise to hexaploid HWG (St1St1St2St2HH).

### Genomic and Evolutionary Drivers of Subgenome Expression Bias

To characterize the expression patterns across subgenomes, we identified 6742 (1:1:1 in H:St_1_:St_2_) homoeologous gene triads across the three subgenomes using synteny mapping. RNA-seq analysis was performed across four tissues—flower, stem, leaf, and root—with three biological replicates per tissue. Principal component analysis (PCA) revealed clear separation of samples by both tissue type and subgenome, with PC1 (30.4%) and PC2 (29.50%) accounting for most of the variance, highlighting distinct expression profiles (Supplementary Figure 13a). Multidimensional Scaling (MDS) further corroborated this tissue-specific clustering (Supplementary Figure 13b). Based on expression patterns across four tissues, we identified 13 distinct co-expression groups among 6742 homologous genes. These clusters reveal the varied expression dynamics of the HWG genome across specific tissues (Supplementary Figure 18).

To analyze the relative contribution of each subgenome to the overall gene expression, we performed comparisons of pairwise gene expression fold changes among subgenomes. The results revealed that the H subgenome harbors more up-regulated genes than either the St1 or St2, whereas St1 and St2 exhibited comparable numbers of up-regulated genes (Figure 4d). This pattern was observed across all four tissues examined (Supplementary Figure 17). In addition, expression divergence between homoeologs was modest, with mean log_2_(*H*/*St_1_*) and log_2_(*H*/*St_2_*) ratios ranging from 0.04 to 0.08 across tissues (Figure 4e), suggesting a weak but detectable H-biased expression. To further investigate homoeologous gene expression bias (HEB), we categorized homoeologous triad into balanced, dominant, and suppressed classes based on the relative expression contributions. Balanced triads predominated in all tissues, accounting from 68.70% to 75.73% of total triads (Supplementary Figure 14). Interestingly, suppressed homoeologs were more frequent than dominant ones across all subgenomes (H: 7.37% vs. 3.95%; St1: 6.10% vs. 1.82%; St2: 6.55% vs. 2.05%; Figure 4a). Notably, the H subgenome showed elevated proportions of both dominant and suppressed homoeologs relative to the St1 and St2 subgenomes. In contrast, the two St subgenomes showed more balanced expression profiles, suggesting greater expression stability between them.

**Fig. 4.**
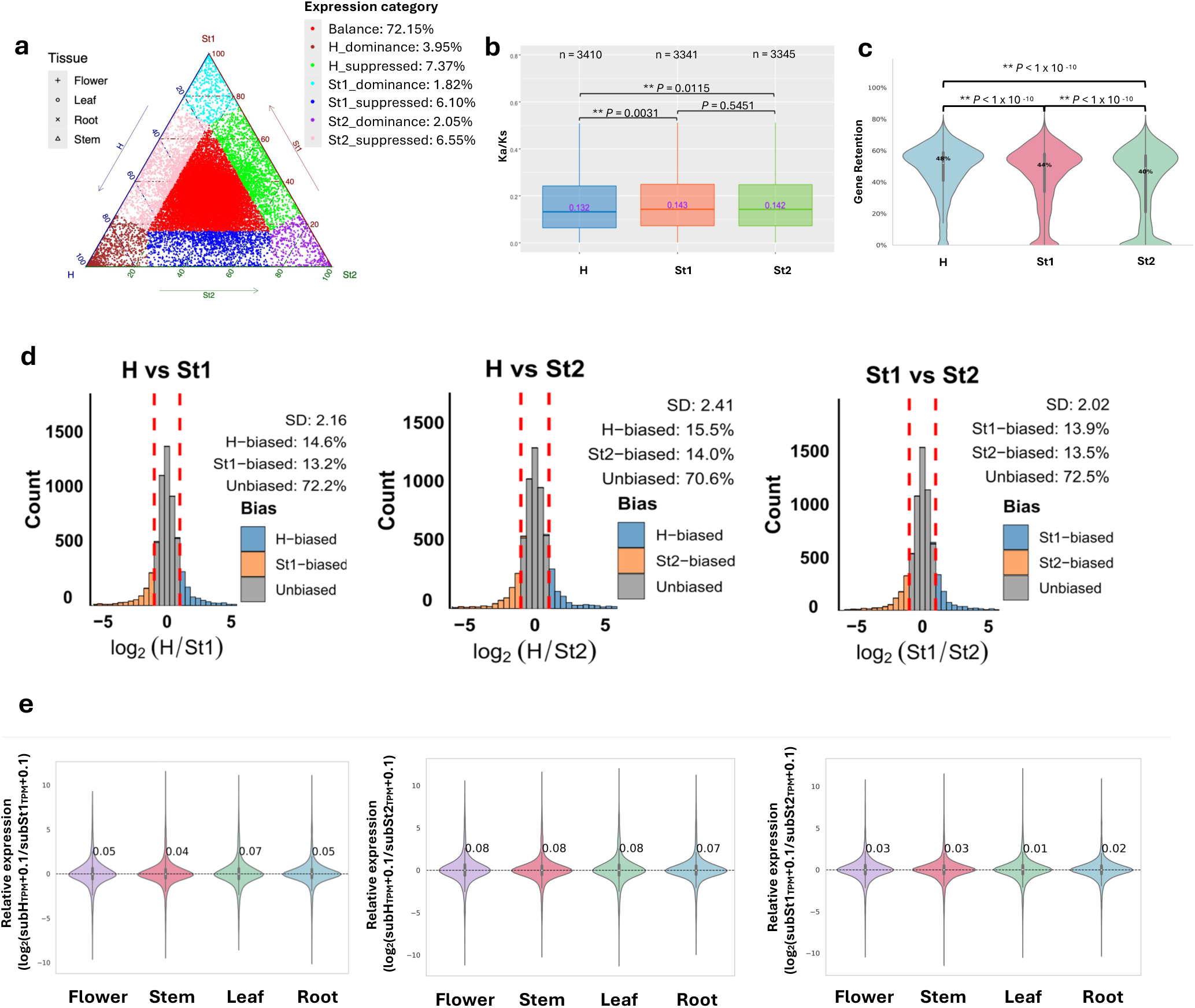
Homoeolog expression patterns in polyploid HWG. **a.** Ternary plot of homoeologs for expression bias across the three subgenome of HWG. Each point in the ternary plot represents a gene triad with coordinates corresponding to the H, St_1_ and St_2_ subgenomes. Triads in vertices indicate dominant categories, whereas triads near edges or between vertices are suppressed categories; balanced triads shown in the center in red. **b**. Box plots comparing *Ka*/*Ks* value distributions among the three subgenomes of HWG. The central line for each box plot denotes the median. Sample size (n) used in each comparison are indicated. The asterisks denote significant differences based on two-tailed Wilcoxon rank-sum test. **c**. Gene retention patterns among subgenomes of HWG relative to the barley genome. The asterisks represent significant differences (based on two-tailed Wilcoxon rank-sum test). **d**. Histograms showing the averaged number of up-regulated genes across four tissues in HWG in each subgenome. **e**. Relative expression of homoeologous gene pairs in four different tissue types.

Next, we investigated possible factors that may contribute to the observed HEB. Long terminal repeats are known regulatory elements that can influence gene expression^30^, therefore, we examined the impact of LTR insertions on protein-coding genes across the subgenomes. Although only minor difference in overall LTR coverage were observed, the flanking regions of genes in the H subgenome showed significantly higher LTR coverage than those in St1 and St2 subgenomes (Supplementary Figure 15a). LTR density is typically negatively associated with subgenome dominance^31,32^, however, this expected relationship was not observed here, suggesting that additional mechanisms contribute to subgenome dominance in HWG. In addition, we assessed gene retention and nonsynonymous-to-synonymous substitution rate ratios (*Ka*/*Ks*) among homoelogs across subgenomes. The H subgenome exhibited a reduced *Ka*/*Ks* ratio implying an accelerated evolutionary rate relative to the St subgenomes, likely reflecting stronger selective pressure. Moreover, the H subgenome retained significantly more genes than St1 and St2 subgenomes (H:48%, St_1_:44%, St_2_:40%) (Figure 4c), suggesting a lower degree of fractionation. This observation implies that that the H genome integrated into the HWG more recently than the St genomes, further supporting our refined evolutionary model. Collectively, these results support the presence of pronounced subgenome-biased gene expression in hexaploid HWG.

### Population Structure and Asymmetric Gene Flow in the Wheatgrass Complex

HWG, originally identified in Central Europe^15^, is an outcrossing polyploid species that has likely undergone extensive reticulate evolution driven by persistent gene flow. Understanding the evolutionary relationships among its close relatives—particularly within the *Elymus* and *Pseudoroegneria* genera—is therefore essential for clarifying how genetic variation is exchanged across their global distributions. However, previous studies have often struggled to clearly delineate the genetic boundaries within and between these groups^33^.

To characterize population structure and resolve these genetic boundaries, we performed GBS on 189 globally sourced accessions, including 71 *E. repens*, 115 *Pseudoroegneria*, 1 *E. dahuricus*, and 2 HWG accessions (Figure 5a). By mapping GBS reads to the HWG reference genome, we identified 50,118 high-quality SNPs. Principal component analysis (Figure 5c) and phylogenetic reconstruction (Figure 5d) consistently resolved three major clades corresponding to *E. repens*, *P. spicata*, and European *Pseudoroegneria* species, in close agreement with their established evolutionary relationships.

**Fig. 5.**
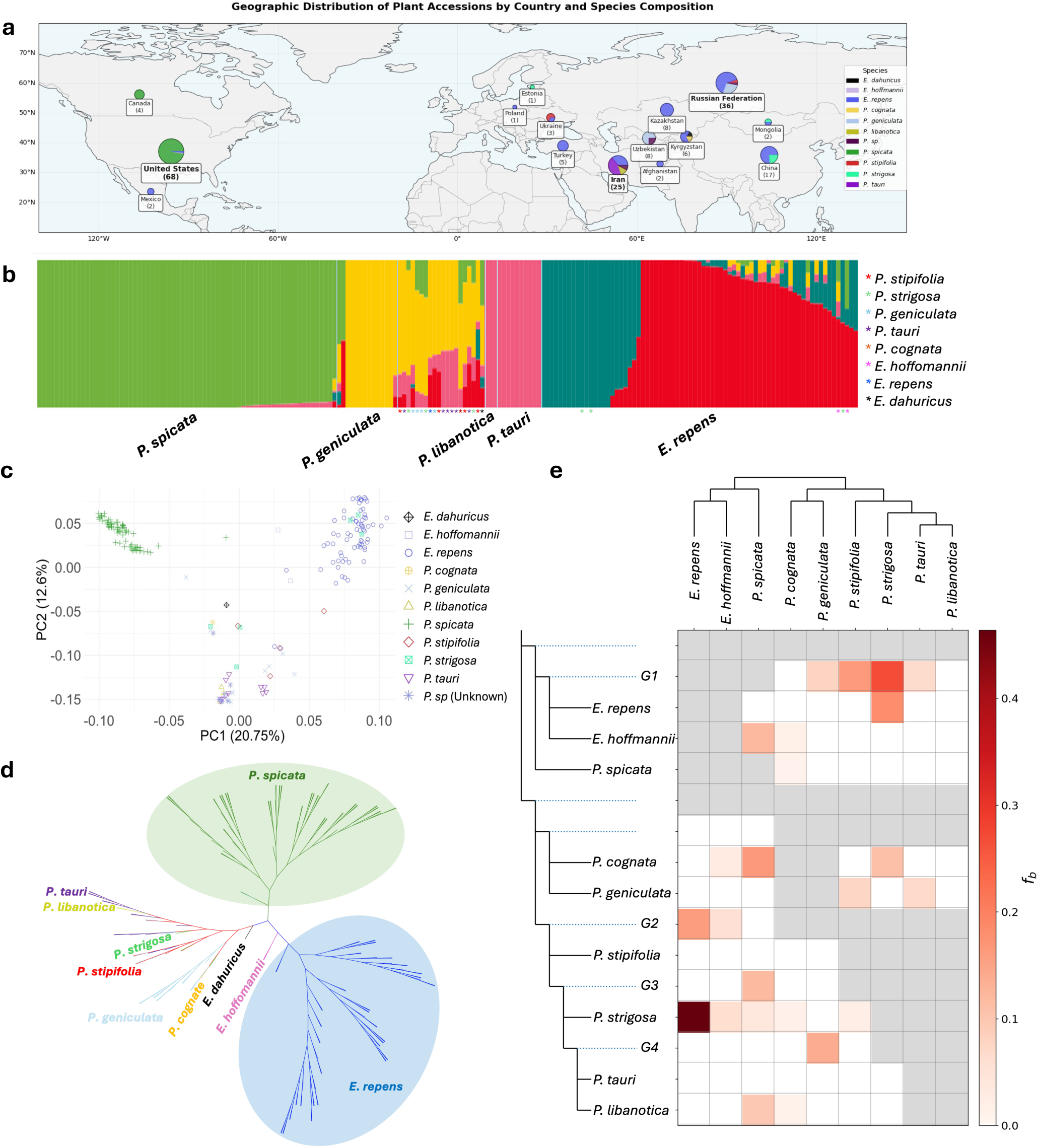
Population structure and interspecific gene flow in the *Elymus* and *Pseudoroegneria* species. **a**, The geographic distribution of collected accessions. Pie charts indicates the proportion of each species in the same country. **b** and **c**, Population structure and principal component analysis (PCA) of 189 accessions including *Pseudoroegneria* and *Elymus* species differentiates *P. spicata*, *E. hoffmannii*, and *E. repens* from other European *Pseudoroegneria* species. Individuals are color coded based on the taxonomic identification. **d**, Dendrogram shows genetic relationship among *Pseudoroegneria* and *Elymus* species. **e**, ABBA-BABA analysis of gene flow within *Pseudoroegneria* and *Elymus* species. Significant introgression events are detected between *E. repens* and *P. strigosa.* An asymmetric pattern of introgression involving *P. spicata*, *P. cognata*, *P. strigosa*, *P. libanotica* and HWG was observed suggesting a significant genetic donor of *P. spicata* to other lineages. G represents groups including multiple species (Methods).

Admixture analysis supported *K*=5 as the optimal number of genetic clusters (lowest cross-validation error: 0.35187). These clusters clearly distinguished *P. spicata*, *P. geniculata*, and *P. tauri*, while subdividing *E. repens* into two distinct subclusters (Figure 5b). Both HWG accessions grouped within the *E. repens* group. Notably, one intermediate cluster spanning *P. geniculata* and *P. tauri* consisted of multiple European *Pseudoroegneria* species (including *P. stipifolia*, *P. strigosa*, *P. tauri*, and *P. cognata*) together with *E. dahuricus*, indicating a shared, highly admixed or hybrid genetic background among these lineages.

To quantify the extent and direction of introgression underlying these patterns, we conducted an *f*-branch (*f_b_*) test across the phylogeny. This analysis revealed strong evidence of gene flow, most prominently between *E. repens* and *P. strigosa* (*f_b_* = 0.464), with weaker but detectable signals between the European clade and the *E. repens*, HWG, and *P. spicata* group (G1) (Figure 5e). A striking feature of the *f*-branch statistic results was the asymmetric role of *P. spicata*. Although it showed little evidence of receiving introgression, *P. spicata* emerged as a major genetic donor to several lineages including HWG (*f_b_* = 0.121), *P. cognata* (*f_b_* = 0.165), *P. strigosa* (*f_b_* = 0.036), and *P. libanotica* (*f_b_* = 0.096) (Figure 5e). This pronounced unidirectional gene flow suggests that *P. spicata* may have served as a key ancestral genetic source during the diversification of these species. Collectively, these results suggest that introgression from *P. spicata* has played a substantial role in shaping the genomic architecture and speciation of European *Pseudoroegneria* lineages.

## Discussion

Polyploid grass genomes are exceptionally large and complex, requiring extensive effort to resolve their genomic structure and evolutionary trajectories^34^. Although chromosome-scale assemblies have become increasingly available across the grass family, particularly for allopolyploids such as oat and wheat^10,35,36^, near-complete assembly of partially autopolyploid species remains rare due to the high sequence similarity among homologous chromosomes. In this study, we generated chromosome-level, haplotype-resolved genome assemblies for CDC Saltking and its diploid ancestors *P. spicata*. These assemblies provide a robust foundation for dissecting subgenome-specific architecture, interspecific variation and evolutionary history within Triticeae.

For decades, the St subgenomes of hexaploid HWG were interpreted as autopolyploid derivatives based on their high sequence similarity and the assumption of genome duplication within a single lineage^15^. Our assembly challenges this simplified view and unveils a more dynamic evolutionary history. While three St_1_ chromosomes are nearly indistinguishable from a canonical St genome, the remaining four chromosomes coalesce into a distinct cluster, indicating unrecognized subgenome differentiation. Most strikingly, Chr4 of the St_2_ subgenome exhibits a unique duplication divergent from St_1_ counterparts in both collinearity and phylogeny (Figure 1d). Comparative genomic analyses demonstrate that this chromosome represents a complex rearrangement incorporating segments from both *P. spicata* and *E. repens* (Figure 1e), implying a large-scale chromosomal recombination or fusion event. Such chromosomal restructuring events are major drivers of karyotype evolution and speciation^37,38^.

Our phylogenomic analyses further refine the origin of the H subgenome. Rather than clustering with barley, the H subgenome of HWG is positioned phylogenetically as an intermediate between *H. marinum* and *H. brevisubulatum*, two species associated with Old-World *Hordeum* clades^39,40^. This finding both corroborates and extends prior observations from *E. repens*, where the H genome was shown to affiliate with *H. brevisubulatum* and *H. bogdanii*, highlighting repeated contributions from ecologically distinct Old-World *Hordeum* lineages^41,42^. Tracing the H genome’s ancestry offers a strategic blueprint for accelerating the development of salt-tolerant crops. Given the close phylogenetic relationship of H genome of HWG with *H. marinum and H. brevisubulatum*, the pronounced salt tolerance observed in HWG likely reflects the retention or parallel evolution of stress-adaptive traits characteristic of *H. marinum*^39^ and *H. brevisubulatum*^40^, two species well known for their ability to thrive under high salinity. This evolutionary proximity strengthens the biological significance of HWG’s salt tolerance, highlighting its role of a valuable reservoir of salinity-adaptive alleles within the Triticeae, with potential relevance for improving salt tolerance in related cereal and forage crops.

Moreover, the functional importance of the H subgenome is further evidenced by its clear expression dominance. Interestingly, this dominance is not accompanied by reduced LTR density. Instead, this suggests that the H subgenome, as the most recent addition to the HWG complex, may be subject to strong positive selection to rapidly integrate into the host genomic background. Such selective pressure could favor the retention and upregulation of functionally important H-derived genes, enhancing overall fitness and stabilizing the hybrid genome.

Interspecific introgression is a recognized driver of adaptive evolution in forage grasses, including perennial ryegrass (*Lolium perenne L.*)^43^, turtle grass (*Thalassia testudinum*)^44,45^, and turfgrass (*Poa trivialis L.*)^46^. Recent phylogenetic studies have proposed that *P. spicata* is originated from a small founder population in North America and may have evolved as a more ancient lineage rather than *P. libanotica*^33^. Here, our *f_b_* statistic analysis supports this evolutionary framework, identifying *P. spicata* as a primary genetic donor to multiple lineages, including HWG, *P. cognata*, *P. strigosa*, and *P. libanotica*. These findings reinforce the hypothesis that *P. spicata* acted as an ancestral source of gene flow that facilitated the speciation of European lineages. Furthermore, the robust signal of introgression between *E. repens* and *P. strigosa*—when compared to the gene flow between HWG and *P. spicata*—suggests that differential introgression events are the primary drivers of the genomic divergence observed between HWG and *E. repens*.

In conclusion, the discovery of differentiated St subgenomes shaped by chromosomal rearrangement, together with evidence for an Old-World *Hordeum* origin of the H subgenome, fundamentally revises prevailing models of hexaploid HWG evolution. These findings underscore the importance of high-resolution, haplotype-resolved assemblies for disentangling complex polyploid histories. Future research integrating high-resolution cytogenetics, transcriptomics, and functional genomics will be instrumental for fully elucidating the evolutionary and agronomic significance of these genomic innovations.

## Material and Methods

### Plant materials and genome sequencing

Hybrid wheatgrass (*E. hoffmannii*, common name:CDC Saltking) was sourced from Dr. Bill Biligetu’s lab at the University of Saskatchewan, Canada. *P. spicata* (PI635993) was sourced from USDA-ARS National Plant Germplasm System in Ames, Iowa, USA. Genome size and ploidy were confirmed through flow cytometry using *H. vulgare* cv. Morex (∼ 5.04 Gb) and Durum wheat strongfield (∼ 12 Gb) as reference genome. DNA was extracted from 8-week-old leaves using NucleoBond HMW DNA kit (Macherey-Nagel). For HWG, DNA libraries were sequenced on a PacBio Revio platform, generating 636.51 Gb HiFi reads (∼ 57 × coverage). In addition, 350 bp DNA-insert libraries were sequenced using an Illumina Novaseq 6000 platform (paired-end 150 bp reads), generating 191.55 Gb paired-end short reads (∼ 17 × coverage). Eight Hi-C libraries were constructed from cross-linked chromatins using a Dovetail Omni-C kit (Dovetail genomics) and sequenced on the Illumina Novaseq 6000 platform, yielding 920.55 Gb Hi-C reads (∼ 82 × coverage). For *P. spicata*, Oxford Nanopore Technologies (ONT) libraries were prepared using Ligation sequencing kit V14 (SQK-LSK114). The ONT system with PromethION flow cells generated 146.63 Gb long reads (∼ 42 × coverage). Using the same Hi-C technology as HWG, 91.55 Gb Hi-C reads (∼ 25 × coverage) were generated for scaffolding.

### Genome size estimation

The genome size of HWG, *P. spicata*, and *E. repens* were estimated through flow cytometry. Chopped fresh shoots from six-week plants were incubated in 1 mL of nuclei extraction buffer (CyStain UV Precise P Automate from Sysmex) for 30 min to release nuclei. The nuclei were collected by filtration through a 40 um of cell strainer, followed by the addition of equal volume of DAPI-based staining buffer (CyStain UV Precise P Automate from Sysmex) and incubated on ice in the dark for 30 min. The fluorescence intensity of DAPI-stained nuclei was quantified using the flow cytometer CytoFLEX (Beckman Coulter, USA). Three replicates were analyzed for each sample. For *P. spicata*, *H. vulgare* (1C = 5 Gb) was used as reference^47^. For HWG and *E. repens*, Durum wheat strongfield cultivar (1C = 12 Gb) was used as reference^48^. Fluorescence histograms were produced with CytExpert software.

### Genome assembly and annotation

For HWG, contig-level assembly was performed using hifiasm v 0.20.0^49^ with default parameters taking HiFi reads as input. For *P. spicata*, contig-level assembly was conducted using hifiasm v0.25.0 with –ont parameter including self-correction and only ONT reads as input^50^. Hi-C scaffolding for both species was conducted by HapHiC^20^ pipelines. Contigs were scaffolded through a four-step process—clustering, reassignment, ordering and orientation, and scaffold construction—followed by manual curation. The Hi-C contact matrix was visualized and manually corrected by Juicebox (v2.20.00)^51^.

Gene structures were predicted integrating *ab initio* prediction with Helixer v0.3.5^52^ (https://github.com/weberlab-hhu/Helixer) and transcriptome data based prediction with Braker3 v.3.0.8^53^. Total RNA was extracted from four different tissues using RNeasy Plant Mini Kit (Qiagen) to construct the RNA-seq library. RNA-seq libraries were prepared using Illumina stranded mRNA Prep Kit (Illumina). RNA-seq was sequenced in Novaseq 6000 platform, generating 300.35 Gb. The gene prediction results from Helixer, Braker 3 were integrated into a final gene annotation set using EVidenceModeler (v1.1.1)^54^. HWG and *P. spicata* followed a similar pipeline for gene annotation.

### TE annotation

To investigate the transposable element landscape in assembled genomes, we used EDTA v.2.1.0^55^ to identify and annotate TEs in the HWG and *P. spicata*. The TE libraries were further classified using TEsorter^56^ and annotated into specific clades.

### Quality assessment of genome assembly and gene annotation

The quality of the assembled genomes is evaluated by LTR assembly index (LAI). The LAI was calculated for hap1 of HWG and *P. spicata* using intact LTR-RTs identified by LTR_FINDER v.1.07^57^ and LRTharvest v.1.1^58^ in EDTA v2.2 pipeline^55^. To quantify the completeness of genomic data and gene annotation, the BUSCO quality assessment has been performed for the assembled genomes and protein sequences with poales_odb12 database using BUSCO v.5.8.2^59^.

### Centromeric region identification

To localize centromeric regions across the assembled genomes, we integrated multiple diagnostic markers and repeat density profiles into a consolidated evidence-based approach. We utilized the BAC7 (AY040832) sequence, previously validated via *in situ* hybridization as a centromere-specific marker in barley^60^. Alongside the distribution of centromeric retrotransposon of maize (CRM) elements, which are highly conserved and typically concentrated within the centromeric cores of grass species^61,62^. In addition, we incorporated three satellite repeats (STlib_96: 528 bp, STlib_98: 503 bp, STlib_117:352 bp), known for their enrichment in the centromeric regions of diploid *P. libanotica*. Notably, STlib_96 and STlib_98 are specific to the St genome and absent in wheat^63^. The density of each marker was calculated across the genome using a 1Mb sliding windows, with centromeric regions defined by the co-localization and significant enrichment of these markers.

### SNP and InDel detection for heterozygosity estimation

Trimmomatic v.0.39 program^64^ was used to trim the adapters, short reads and poor-quality data for raw Illumina reads of HWG. A total of 2,129,824,780 pair-end reads for HWG passed filtering and were retained in downstream analysis. High quality reads were aligned to the assembled genome of HWG using Burrows-Wheeler Aligner (BWA-MEM) v 0.7.18 to generate bam files^65^. GATK (v4.3.0)^66^ variant calling pipeline was used to call SNPs and INDELs and only high-quality variants (MQ > 30) are maintained in the final VCF files^66^. The heterozygosity and SNP density were calculated based on the VCF file.

### Subgenome assignment

To differentiate the chromosomes between subgenomes, we identified the differential *k*-mers among homoeologous chromosome sets in HWG (hap1) using SubPhaser^23^. This strategy has been successfully applied to resolve subgenomes in several polyploid plant species^9,67^. Heterogenous chromosomes were clustered based on the enriched *k*-mers by a *k*-means algorithm. To verify the accuracy of the results, we have also tested different *k*-mer and minimum counts combinations (*k* = 15, 30, 33 and q = 50, 100, 300), which produced fully consistent clustering pattern with significant higher number of differential *k*-mers in H subgenome than two St subgenomes. Finally, we identified 690,497 differential *k*-mers in HWG (*k* = 15 and frequency ≥ 50). The H subgenome contributed the majority of these *k*-mers (631,056), whereas St1 and St2 contributed 23,871 and 35,569 *k*-mers, respectively. To characterize the identity of chromosomes in each clustering, we performed the dotplot analysis to identify the exact matching regions between the HWG and two reference genomes (*P. spicata* and *E. sibiricus*) using MUMMER v.3.23 (nucmer -c 100 -l 200/300)^68^. To differentiate between conserved and diverged subgenomic regions, we systematically varied the minimum exact length in the alignment parameters. A stringent threshold of 300 bp was utilized to identify highly conserved chromosome regions, while a reduced threshold of 200 bp was applied to capture more diverged homoeologous segments, specifically to resolve chromosomes (Chr3, Chr4, and Chr6) in St_1_ subgenome that were potentially misclassified during *k*-mer analysis.

To further verify the homologous chromosome pairs, genome-wide synteny analysis was performed using Barley as a reference. The syntenic blocks were identified using WGDI (Whole-Genome Duplication Integrated analysis, v0.74) software package with the -a alignment function^69^.

### Phylogenetic analysis and divergence time estimation

A total of 975 single-copy genes conserved in land plants from 11 Triticeae species (*Oryza sativa*^70^, *Brachypodium distachyon*^71^, *Hordeum vulgare*^47^, *Hordeum spontaneum*^72^, *Hordeum marinum*^39^, *E. sibiricus*^22^, *Secale cereale*^73^, *Thinopyrum elongatum*^21^, wheat^74^, HWG, and *P. spicata*) including 16 subgenomes were identified by OrthoFinder^75^. The protein sequence alignments were first generated using Clustalw (v.1.83)^76^ in a fasta format. For each single-copy gene, protein trees were inference using IQ-TREE2 v 2.2.03^77^ with LG model. The 975 single-copy gene trees were concatenated and subjected to the species tree inference with MFP model. *Oryza sativa* was used as the outgroup for rooting. Gene tree discordance within 975 genes was quantified and visualized by pie charts for the nodes related to HWG.

MCMCTREE in the PAML (v4.9) package^78^ was used to estimate divergence times with default parameters in phylogenetic analysis. The divergence time of *Oryza sativa* (47.85 Mya) was used for calibration^10^.

The chloroplast genomes from HWG and *P. spicata* were *de novo* assembled using Oatk software^79^ (-k 1001 -c 150) with HiFi reads (HWG) and corrected ONT Ultra-long reads (*P. spicata*), respectively. The chloroplast genome sequences for the rest species were downloaded from NCBI (*Hordeum vulgare*: NC_056985.1, *Brachypodium distachyon*: NC_011032.1, *Thinopyrum elongatum*: NC_043841.1, *Elymus sibiricus*: NC_058919.1, *Hordeum marinum*: OR063947.1, *Oryza sativa*:NC_031333.1, *Secale cereale*: NC_021761.1, *Triticum aestivum*: NC_002762.1). With *Oryza sativa* as outgroup, the 10 plastome sequences were first aligned using Clustalw (v.1.83) and subsequently trimmed using trimAl (v1.4)^80^ with default parameters. A maternal phylogenetic tree was inferred using IQ-TREE2 with LG model with 1000 bootstrap.

### Phylogenetic tree construction using SNPs

For the phylogenetic analysis in Figure 3f, Illumina reads from all species were initially mapped to the HWG reference genome. Unique reads specifically mapping to each subgenome were isolated and re-mapped to their respective subgenomes for independent SNP calling via the GATK pipeline^66^. The resulting SNP sets (VCF format) were converted into aligned FASTA format using vcf2phylip^81^. Finally, a maximum-likelihood tree was inferred using IQ-TREE2 v2.2.03^77^, utilizing the aligned FASTA as input and the best-fit model selection (-m AUTO).

### *K*a/*K*s analysis

3,422 one-to-one orthologous gene sets were fetched from OrthoFinder results for the 8 subgenomes in 5 species (HWG, *P. spicata*, *E. sibiricus*, *H. vulgare*, *H. marinum*). The protein sequences from each gene pair were aligned using MUSCLE^82^ and subsequently converted to a nucleotide alignment by using PAL2NAL (v14)^83^. The *K*a and *K*s values were computed between the gene pairs using Codeml from PAML (v4.9)^84^. To minimize the noise from extremely low *K*s values, estimates that lower than 0.01 were removed for histogram. The *K*s values of all gene pairs were used to determine the divergence time of subgenomes. For the *K*a/*K*s analysis, all the *K*a and *K*s values are compared to the corresponding genes in *H. vulgare*. The *K*a/*K*s ratio for all genes in each subgenome were used to estimate the selective pressure for each subgenome. The significance of the differences in *K*a/*K*s ratios between subgenomes was estimated based on Wilcoxon rank-sum test for non-normal distribution in R.

### Collinearity analysis among species

Collinearity among the subgenomes of HWG, *H. vulgare*, *H. marinum*, *E. sibiricus*, and *P. spicata* was individually analyzed for each pair using JCVI v.1.5.6 software package^85^. Homoeologous gene pairs were first identified based on gene comparison by LAST. Genes exhibiting best hits were maintained and considered as homoeologs. Syntenic blocks were identified through single linkage clustering performed on LAST output with default settings (jcvi.compara.synteny function). The syntenic blocks shared between the H and St subgenomes from different species were visualized with the jcvi.graphics.karyotype function.

### Homoeologous expression bias (HEB) analysis

RNA was isolated from four types of tissues from HWG, including flower, stem, leaf, and root using RNeasy Plant Mini Kit (Qiagen) to construct the RNA-seq library. The paired-end reads from RNA-seq were subjected to quality trimming using Trimmomatic (v0.39) with the default settings and aligned to the HWG genome with STAR (v2.7.6) software^86^ with default parameters. Gene expression level was quantified using RSEM (v1.3.3) program^87^. The expression level in transcripts per million (TPM) values were used in expression analysis.

Homoeologous gene triplets were initially identified based on the syntenic blocks detected using JCVI (v.1.5.6). Protein sequences within each homoeologous group were aligned using Clustalw (v.1.83) to verify sequence similarity and confirm homology. A total of 6742 gene triplets (triads) in hexaploid HWG were ultimately selected for differential homoeologous gene expression analysis (Supplementary Table 3). To cluster the homoeologous genes, TPM values of triplets across all tissues were transformed using the common logarithm. A Euclidean distance matrix was computed from normalized data, followed by two-dimensional hierarchical clustering to group homoeologous genes with similar expression profiles. To evaluate the relative expression of homoeologous gene pairs in different tissues, the relative expression between two subgenomes was calculated as the log base 2 of the ratio between the TPM value of each subgenomes with applying a pseudocount to avoid division by zero (log_2_(Sub_1TPM_ + 0.1/Sub_2TPM_ + 0.1).

To analyze the subgenome bias expression in HWG, we normalized the absolute TPM for each gene within a homoeologous triad by calculating its normalized expression as the proportion of its TPM value divided by the sum of TPM values from all three subgenomes. To improve the reliability of subgenome bias analysis and avoid artificial bias, the triplets with the sum of TPM value across the three subgenomes less than 0.5 were removed. Based on the normalized expression proportions of the three homoeologs within each triad, genes were classified into expression bias categories. Triads with proportions close to *1:1:1* were considered balanced. A proportion close to 1:0:0 indicated H-subgenome dominance, while 0:1:0 and 0:0:1 represented St_1_-subgenome dominance and St_2_-subgenome dominance, respectively. Conversely, proportions such as 0:0.5:0.5 indicated H-subgenome suppression, whereas 0.5:0:0.5 and 0.5:0.5:0 indicated St_1_- and St_2_-subgenome suppression, respectively. The normalized expression analysis was performed for each tissue as well as the average across all tissues.

### Gene retention analysis

To investigate the effect of gene retention on HEB, we estimated the retention rates of conserved genes in syntenic blocks in each subgenome of HWG using *H. vulgare* as reference (Supplementary Figure 16). For this purpose, the “retain” function (wgdi -r) in the Python-based tool WGDI (v0.74) has been used to assess the gene retention^69^.

### LTR coverage estimation in the flanking regions of genes

For all the genes in each subgenome, 5 kb sequences in upstream (5’) and downstream (3’) were extracted as flanking regions. BEDTools (v2.30.0)^88^ program (makewindows sub-command) was used to generate 100 bp sliding windows with a 10 bp step across these flanking regions. For each window, the coverage of LTR sequences was quantified with the sub-command intersect -c in BEDTools, and the average of coverage was subsequently calculated for all genes in each subgenome.

### GBS sequencing and data analysis

Genotyping-by-sequencing libraries was prepared based on the method described by Ji *et al*. (2025)^33^. Briefly, 200 ng of DNA from each sample were digested with the CHG methylation sensitive restriction enzymes *PstI* and *MspI* followed by indexing and PCR amplification. After PCR, the library was purified and quantified by Bioanalyzer (Agilent Technologies) to confirm the fragment size and quality. The libraries were sequenced on a NovaSeq6000 platform at OPAL. Raw reads were demultiplexed using an in-house script. Trimmomatic (v.0.39)^64^ program was used to trim the adapters, and remove the short reads and poor-quality data. The cleaned reads were mapped to the HWG reference genome using BWA v0.7.18^65^. Variants from 189 accessions were called using BCFtools^89^. A total of 206,132,162 variants including SNPs and Indels were identified. The Bi-allelic SNPs were first isolated from the variants file using SAMtools and BCFtools^65,90^. The redundant SNPs were removed using a linkage disequilibrium pruning procedure with PLINK^91,92^, with a window size of 50 kb, step size of 10 SNPs, and *r^2^* threshold of 0.2. The core SNPs were further filtered with > 10% missing rate, MAF < 0.05, and Hardy-Weinberg Equilibrium p-value < 1e-6. The final number of SNPs after filtration is 50,118.

### Population structure analysis and phylogenetic dendrogram construction

We determined the choice of *k* using a 5-fold cross-validation (CV) procedure. A number of ancestral populations (*k* value) ranging from 2 to 9 were tested. The optimal *k* value (*k* = 5) was determined based on the lowest CV error indicating that five ancestral genetic clusters best explain the population structure in our case. Population structure was analyzed using ADMIXTURE v1.3.0 program^93^. The corresponding Q matrix from *k* = 5 model was computed and visualized by TBtools (v2.371)^94^ to illustrate the admixture patterns among populations. For phylogenetic tree construction, the filtered SNPs dataset including all the accessions in VCF format was converted to aligned fasta format by using vcf2phylip^81^. A maximum likelihood tree was inferred by IQ-TREE2 v 2.2.03^77^ with best-fit model (-m AUTO) using aligned fasta as input.

### ABBA-BAAB test

On the basis of SNPs variants identified from 189 accessions, an ABBA-BABA test (or D-statistic) was performed between all possible triplets among lineages setting *Elymus dahuricus* as the outgroup using Dsuite software (v.0.4 r28)^95^. Dtrios command was used to calculate the D statistic and the *f*4-ratio for all combinations of trios of species with -t parameter. The *f*-branch metric was calculated with subcommand Fbranch indicating the gene flow between specific branches including individual species and groups. Four designated groups (G1 to G4) within the introgression matrix are: G1 (*P. spicata, E. repens, E. hoffmannii*), G2 (*P. stipifolia, P. strigosa, P. tauri, P. libanotica*), G3 (*P. strigosa, P. tauri*, *P. libanotica*), G4 (*P. tauri, P. libanotica*). Given the complex genetic signals and potential reticulation identified in the line-level analysis, we inferred a consensus species-level tree based on the placement of the majority of accessions. This tree topology serves as the proposed evolutionary framework for gene flow signal visualization.

## Data availability

The genome sequence data of both haplotypes of HWG and *P. spicata* are deposited in NCBI with BioSample accessions: SAMN56685049, SAMN56685050, SAMN56685051, SAMN56685052 under BioProject PRJNA1440508. The WGS and GBS datasets are deposited in NCBI under BioProject PRJNA1440508.

## Code availability

The custom codes in this study are available at Github (https://github.com/USask-BINFO/Hybrid_wheatgrass_genome.git)

## Acknowledgements

We thank Wentao Zhang from National Research Council Canada for providing the Durum wheat seeds for flow cytometry analysis. We also extend our gratitude to the Germplasm Resources Information Network (GRIN) for providing the seeds material for our study, and support from Omics and Precision Analytics Laboratory (OPAL) and Data Management and Analytics (DMA) platform at GIFS for DNA sequencing and analysis. This work is funded by Natural Sciences and Engineering Research Council of Canada, the Agriculture Development Fund (ADF) under the Ministry of Agriculture of Saskatchewan and the Saskatchewan Cattleman’s Association.

## Author contributions

Y.J., L.J. and A.S. conceptualized the study. Y.J., R.C., S.P. and N.K. performed genome assembly and annotation. Z.W. performed Hi-C sequencing and RNA-seq experiments for *P. spicata*. Y.J. and N.K. performed the analysis of subgenome dominance. L.M.M. analyzed Ks distribution. P.H. and B.B. provide HWG and Quackgrass species material. Y.J. worked on all the other experiments and analysis. Y.J., L.J. and A.S. wrote the paper. All authors reviewed and approved the final version of the manuscript.

## Additional information

Supplementary data is available for this paper at: https://github.com/ifoo1213/Wheatgrass_genome/blob/main/Supplementary_tables

## Reference

1. Lesk, C., Rowhani, P. & Ramankutty, N. Influence of extreme weather disasters on global crop production. Nature 529, 84–7 (2016).

2. Lobell, D.B., Schlenker, W. & Costa-Roberts, J. Climate trends and global crop production since 1980. Science 333, 616–20 (2011).

3. Myers, S.S., et al. Climate change and global food systems: potential impacts on food security and undernutrition. Annu Rev Public Health 38, 259–277 (2017).

4. Steppuhn, H. & Asay, K. Emergence, height, and yield of tall, NewHy, and green wheatgrass forage crops grown in saline root zones. Canadian journal of plant science 85, 863–875 (2005).

5. Waldner, A.A. Unpublished master’s thesis, University of Saskatchewan (2025).

6. Jensen, K.B., Peel, M.D., Waldron, B.L., Horton, W.H. & Asay, K.H. Persistence after three cycles of selection in NewHy RS-wheatgrass (*Elymus hoffmannii* KB Jensen & Asay) at increased salinity levels. Crop science 45, 1717–1720 (2005).

7. Capstaff, N.M. & Miller, A.J. Improving the yield and nutritional quality of forage crops. Frontiers in Plant Science 9, 535 (2018).

8. Jungers, J.M., DeHaan, L.R., Betts, K.J., Sheaffer, C.C. & Wyse, D.L. Intermediate wheatgrass grain and forage yield responses to nitrogen fertilization. Agronomy Journal 109, 462–472 (2017).

9. Jin, X., et al. Haplotype-resolved genomes of wild octoploid progenitors illuminate genomic diversifications from wild relatives to cultivated strawberry. Nature Plants 9, 1252–1266 (2023).

10. Peng, Y., et al. Reference genome assemblies reveal the origin and evolution of allohexaploid oat. Nature Genetics 54, 1248–1258 (2022).

11. Zhuang, Y., et al. Phylogenomics of the genus Glycine sheds light on polyploid evolution and life-strategy transition. Nature Plants 8, 233–244 (2022).

12. Steppuhn, H., Jefferson, P., Iwaasa, A. & McLeod, J. AC Saltlander green wheatgrass. Canadian Journal of Plant Science 86, 1161–1164 (2006).

13. Asay, K., et al. Registration of ‘NewHy’ RS hybrid wheatgrass. (1991).

14. Asay, K.H. RS hybrid wheatgrass.

15. Jensen, K. & Asay, K. Cytology and morphology of *Elymus hoffmanni* (Poaceae: Triticeae): a new species from the Erzurum Province of Turkey. International Journal of Plant Sciences 157, 750–758 (1996).

16. Ramsey, J. & Schemske, D.W. Pathways, mechanisms, and rates of polyploid formation in flowering plants. Annual review of ecology and systematics 29, 467–501 (1998).

17. Van de Peer, Y., Mizrachi, E. & Marchal, K. The evolutionary significance of polyploidy. Nature Reviews Genetics 18, 411–424 (2017).

18. Brauer, C.J., et al. Natural hybridization reduces vulnerability to climate change. Nature Climate Change 13, 282–289 (2023).

19. Luong, K.T.M., Georgievich, D.M., Anh, N.P., Vitalievna, K.A. & Iliych, K.G. Differences in ploidy level and genome constitution revealed by cytogenetic analysis of *Pseudoroegneria* germplasm accessions: case study. Известия Тимирязевской сельскохозяйственной академии, 29–35 (2015).

20. Zeng, X., et al. Chromosome-level scaffolding of haplotype-resolved assemblies using Hi-C data without reference genomes. Nature plants 10, 1184–1200 (2024).

21. Wang, H., et al. Horizontal gene transfer of *Fhb7* from fungus underlies Fusarium head blight resistance in wheat. Science 368, eaba5435 (2020).

22. Shen, W., et al. Chromosome-scale assembly of the wild cereal relative *Elymus sibiricus*. Scientific Data 11, 823 (2024).

23. Jia, K.H., et al. SubPhaser: a robust allopolyploid subgenome phasing method based on subgenome-specific *k*-mers. New Phytologist 235, 801–809 (2022).

24. Fuller, Z.L., Koury, S.A., Phadnis, N. & Schaeffer, S.W. How chromosomal rearrangements shape adaptation and speciation: case studies in *Drosophila pseudoobscura* and its sibling species *Drosophila persimilis*. Molecular ecology 28, 1283–1301 (2019).

25. Berdan, E.L., et al. Structural variants and speciation: multiple processes at play. Cold Spring Harbor Perspectives in Biology 16, a041446 (2024).

26. Cheng, H., et al. Frequent intra-and inter-species introgression shapes the landscape of genetic variation in bread wheat. Genome biology 20, 136 (2019).

27. Tyrka, M., et al. Evaluation of genetic structure in European wheat cultivars and advanced breeding lines using high-density genotyping-by-sequencing approach. BMC genomics 22, 81 (2021).

28. Zhang, L., et al. Phylotranscriptomics resolves the phylogeny of Pooideae and uncovers factors for their adaptive evolution. Molecular biology and evolution 39, msac026 (2022).

29. Rosenberg, N.A. & Nordborg, M. Genealogical trees, coalescent theory and the analysis of genetic polymorphisms. Nature Reviews Genetics 3, 380–390 (2002).

30. Bubb, K.L., et al. The regulatory potential of transposable elements in maize. Nature Plants, 1–12 (2025).

31. Zhang, K., et al. The lack of negative association between TE load and subgenome dominance in synthesized *Brassica* allotetraploids. Proceedings of the National Academy of Sciences 120, e2305208120 (2023).

32. Li, C., et al. Extraordinary preservation of gene collinearity over three hundred million years revealed in homosporous lycophytes. Proceedings of the National Academy of Sciences 121, e2312607121 (2024).

33. Ji, Y., et al. Genomic insights into wheatgrass: unravelling genetic diversity, population structure, and evolutionary dynamics in *Pseudoroegneria* species. Molecular Phylogenetics and Evolution, 108397 (2025).

34. Zhang, X., Wu, R., Wang, Y., Yu, J. & Tang, H. Unzipping haplotypes in diploid and polyploid genomes. Computational and structural biotechnology journal 18, 66–72 (2020).

35. He, Q., et al. The near-complete genome assembly of hexaploid wild oat reveals its genome evolution and divergence with cultivated oats. Nature Plants 10, 2062–2078 (2024).

36. Liu, S., et al. A telomere-to-telomere genome assembly coupled with multi-omic data provides insights into the evolution of hexaploid bread wheat. Nature Genetics, 1–13 (2025).

37. Parkin, I.A., et al. Transcriptome and methylome profiling reveals relics of genome dominance in the mesopolyploid *Brassica oleracea*. Genome biology 15, R77 (2014).

38. Liu, S., et al. The *Brassica oleracea* genome reveals the asymmetrical evolution of polyploid genomes. Nature communications 5, 3930 (2014).

39. Kuang, L., et al. The genome and gene editing system of sea barleygrass provide a novel platform for cereal domestication and stress tolerance studies. Plant Communications 3(2022).

40. Feng, H., et al. *Hordeum* I genome unlocks adaptive evolution and genetic potential for crop improvement. Nature plants, 1–15 (2025).

41. Mason-Gamer, R.J. Allohexaploidy, introgression, and the complex phylogenetic history of *Elymus repens* (Poaceae). Molecular Phylogenetics and Evolution 47, 598–611 (2008).

42. Mahelka, V. & Kopecký, D. Gene capture from across the grass family in the allohexaploid *Elymus repens* (L.) Gould (Poaceae, Triticeae) as evidenced by ITS, GBSSI, and molecular cytogenetics. Molecular Biology and Evolution 27, 1370–1390 (2010).

43. Cunliffe, K., et al. Assessment of gene flow using tetraploid genotypes of perennial ryegrass (*Lolium perenne* L.). Australian Journal of Agricultural Research 55, 389–396 (2004).

44. van Dijk, K.j., Bricker, E., van Tussenbroek, B.I. & Waycott, M. Range-wide population genetic structure of the Caribbean marine angiosperm *Thalassia testudinum*. Ecology and Evolution 8, 9478–9490 (2018).

45. Schlueter, M.A. & Guttman, S.I. Gene flow and genetic diversity of turtle grass, *Thalassia testudinum*, banks ex könig, in the lower Florida Keys. Aquatic Botany 61, 147–164 (1998).

46. Brunharo, C.A., Benson, C.W., Huff, D.R. & Lasky, J.R. Chromosome-scale genome assembly of *Poa trivialis* and population genomics reveal widespread gene flow in a cool-season grass seed production system. Plant Direct 8, e575 (2024).

47. Mascher, M., et al. A chromosome conformation capture ordered sequence of the barley genome. Nature 544, 427–433 (2017).

48. Maccaferri, M., et al. Durum wheat genome highlights past domestication signatures and future improvement targets. Nature genetics 51, 885–895 (2019).

49. Cheng, H., Asri, M., Lucas, J., Koren, S. & Li, H. Scalable telomere-to-telomere assembly for diploid and polyploid genomes with double graph. Nature Methods 21, 967–970 (2024).

50. Cheng, H., et al. Efficient near telomere-to-telomere assembly of Nanopore Simplex reads. bioRxiv, 2025.04. 14.648685 (2025).

51. Durand, N.C., et al. Juicebox provides a visualization system for Hi-C contact maps with unlimited zoom. Cell systems 3, 99–101 (2016).

52. Holst, F., et al. Helixer–de novo prediction of primary eukaryotic gene models combining deep learning and a hidden Markov model. BioRxiv, 2023.02. 06.527280 (2023).

53. Gabriel, L., et al. BRAKER3: Fully automated genome annotation using RNA-seq and protein evidence with GeneMark-ETP, AUGUSTUS, and TSEBRA. Genome research 34, 769–777 (2024).

54. Haas, B.J., et al. Automated eukaryotic gene structure annotation using EVidenceModeler and the Program to Assemble Spliced Alignments. Genome biology 9, R7 (2008).

55. Ou, S., et al. Benchmarking transposable element annotation methods for creation of a streamlined, comprehensive pipeline. Genome biology 20, 275 (2019).

56. Zhang, R.-G., et al. TEsorter: an accurate and fast method to classify LTR-retrotransposons in plant genomes. Horticulture Research 9, uhac017 (2022).

57. Xu, Z. & Wang, H. LTR_FINDER: an efficient tool for the prediction of full-length LTR retrotransposons. Nucleic acids research 35, W265–W268 (2007).

58. Ellinghaus, D., Kurtz, S. & Willhoeft, U. LTRharvest, an efficient and flexible software for de novo detection of LTR retrotransposons. BMC bioinformatics 9, 18 (2008).

59. Manni, M., Berkeley, M.R., Seppey, M., Simão, F.A. & Zdobnov, E.M. BUSCO update: novel and streamlined workflows along with broader and deeper phylogenetic coverage for scoring of eukaryotic, prokaryotic, and viral genomes. Molecular biology and evolution 38, 4647–4654 (2021).

60. Hudakova, S., et al. Sequence organization of barley centromeres. Nucleic Acids Research 29, 5029–5035 (2001).

61. Kejnovsky, E., Jedlicka, P., Lexa, M. & Kubat, Z. Factors determining chromosomal localization of transposable elements in plants. Plant Biology 27, 975–989 (2025).

62. Chang, X., et al. High-quality *Gossypium hirsutum* and *Gossypium barbadense* genome assemblies reveal the landscape and evolution of centromeres. Plant communications 5(2024).

63. Wu, D., et al. *Pseudorogneria libanotica* intraspecific genetic polymorphism revealed by fluorescence in situ hybridization with newly identified tandem repeats and wheat single-copy gene probes. International Journal of Molecular Sciences 23, 14818 (2022).

64. Bolger, A.M., Lohse, M. & Usadel, B. Trimmomatic: a flexible trimmer for Illumina sequence data. Bioinformatics 30, 2114–2120 (2014).

65. Li, H. & Durbin, R. Fast and accurate short read alignment with Burrows–Wheeler transform. bioinformatics 25, 1754–1760 (2009).

66. Van der Auwera, G.A., et al. From FastQ data to high-confidence variant calls: the genome analysis toolkit best practices pipeline. Current protocols in bioinformatics 43, 11.10. 1–11.10. 33 (2013).

67. Mitros, T., et al. Genome biology of the paleotetraploid perennial biomass crop *Miscanthus*. Nature communications 11, 5442 (2020).

68. Kurtz, S., et al. Versatile and open software for comparing large genomes. Genome biology 5, R12 (2004).

69. Sun, P., et al. WGDI: a user-friendly toolkit for evolutionary analyses of whole-genome duplications and ancestral karyotypes. Molecular plant 15, 1841–1851 (2022).

70. International Rice Genome Sequencing Project. The map-based sequence of the rice genome. Nature 436, 793–800 (2005).

71. International Brachypodium Initiative. Genome sequencing and analysis of the model grass *Brachypodium distachyon*. Nature 463, 763–768 (2010).

72. Zhang, W., et al. Genome architecture and diverged selection shaping pattern of genomic differentiation in wild barley. Plant Biotechnol J 21, 46–62 (2023).

73. Rabanus-Wallace, M.T., et al. Chromosome-scale genome assembly provides insights into rye biology, evolution and agronomic potential. Nat Genet 53, 564–573 (2021).

74. International Wheat Genome Sequencing Consortium. A chromosome-based draft sequence of the hexaploid bread wheat (*Triticum aestivum*) genome. Science 345, 1251788 (2014).

75. Emms, D.M. & Kelly, S. OrthoFinder: phylogenetic orthology inference for comparative genomics. Genome biology 20, 238 (2019).

76. Larkin, M.A., et al. Clustal W and Clustal X version 2.0. bioinformatics 23, 2947–2948 (2007).

77. Minh, B.Q., et al. IQ-TREE 2: new models and efficient methods for phylogenetic inference in the genomic era. Molecular biology and evolution 37, 1530–1534 (2020).

78. Yang, Z. PAML: a program package for phylogenetic analysis by maximum likelihood. Comput Appl Biosci 13, 555–6 (1997).

79. Zhou, C., et al. Oatk: a de novo assembly tool for complex plant organelle genomes. Genome Biology 26, 235 (2025).

80. Capella-Gutiérrez, S., Silla-Martínez, J.M. & Gabaldón, T. trimAl: a tool for automated alignment trimming in large-scale phylogenetic analyses. Bioinformatics 25, 1972–1973 (2009).

81. Ortiz, E.M. vcf2phylip v1. 5: convert a VCF matrix into several matrix formats for phylogenetic analysis. Zenodo (2019).

82. Edgar, R.C. MUSCLE: a multiple sequence alignment method with reduced time and space complexity. BMC bioinformatics 5, 113 (2004).

83. Suyama, M., Torrents, D. & Bork, P. PAL2NAL: robust conversion of protein sequence alignments into the corresponding codon alignments. Nucleic acids research 34, W609–W612 (2006).

84. Yang, Z. PAML 4: phylogenetic analysis by maximum likelihood. Molecular biology and evolution 24, 1586–1591 (2007).

85. Tang, H. et al. JCVI: A versatile toolkit for comparative genomics analysis. Imeta 3, e211 (2024).

86. Dobin, A., et al. STAR: ultrafast universal RNA-seq aligner. Bioinformatics 29, 15–21 (2013).

87. Li, B. & Dewey, C.N. RSEM: accurate transcript quantification from RNA-Seq data with or without a reference genome. BMC bioinformatics 12, 323 (2011).

88. Quinlan, A.R. & Hall, I.M. BEDTools: a flexible suite of utilities for comparing genomic features. Bioinformatics 26, 841–842 (2010).

89. Danecek, P., et al. Twelve years of SAMtools and BCFtools. Gigascience 10, giab008 (2021).

90. Li, H., et al. The sequence alignment/map format and SAMtools. bioinformatics 25, 2078–2079 (2009).

91. Purcell, S., et al. PLINK: a tool set for whole-genome association and population-based linkage analyses. The American journal of human genetics 81, 559–575 (2007).

92. Chang, C.C., et al. Second-generation PLINK: rising to the challenge of larger and richer datasets. Gigascience 4, s13742-015-0047-8 (2015).

93. Alexander, D.H., Novembre, J. & Lange, K. Fast model-based estimation of ancestry in unrelated individuals. Genome research 19, 1655–1664 (2009).

94. Chen, C., et al. TBtools: an integrative toolkit developed for interactive analyses of big biological data. Molecular plant 13, 1194–1202 (2020).

95. Malinsky, M., Matschiner, M. & Svardal, H. Dsuite-Fast D-statistics and related admixture evidence from VCF files. Molecular ecology resources 21, 584–595 (2021).

