## Supplementary_Figures for "Haplotype-resolved Genome Assemblies Reveal Subgenome Origins, Genome Bias and Reticulate Evolution in Wheatgrass Species"

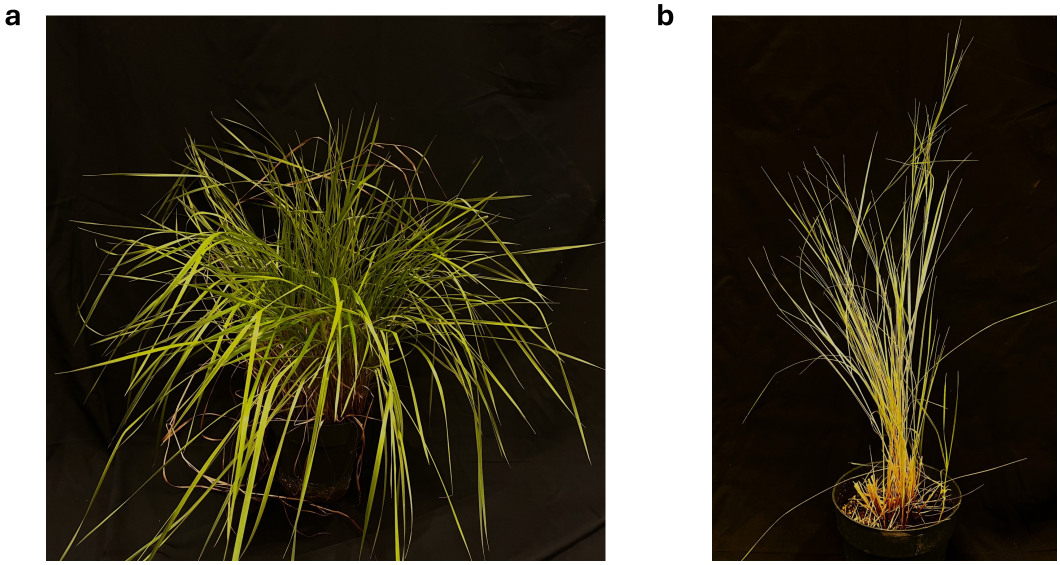


**Supplementary Figure 1| Morphology of the sequenced species *E. hoffmannii* (a) and *P. spicata* (b)**

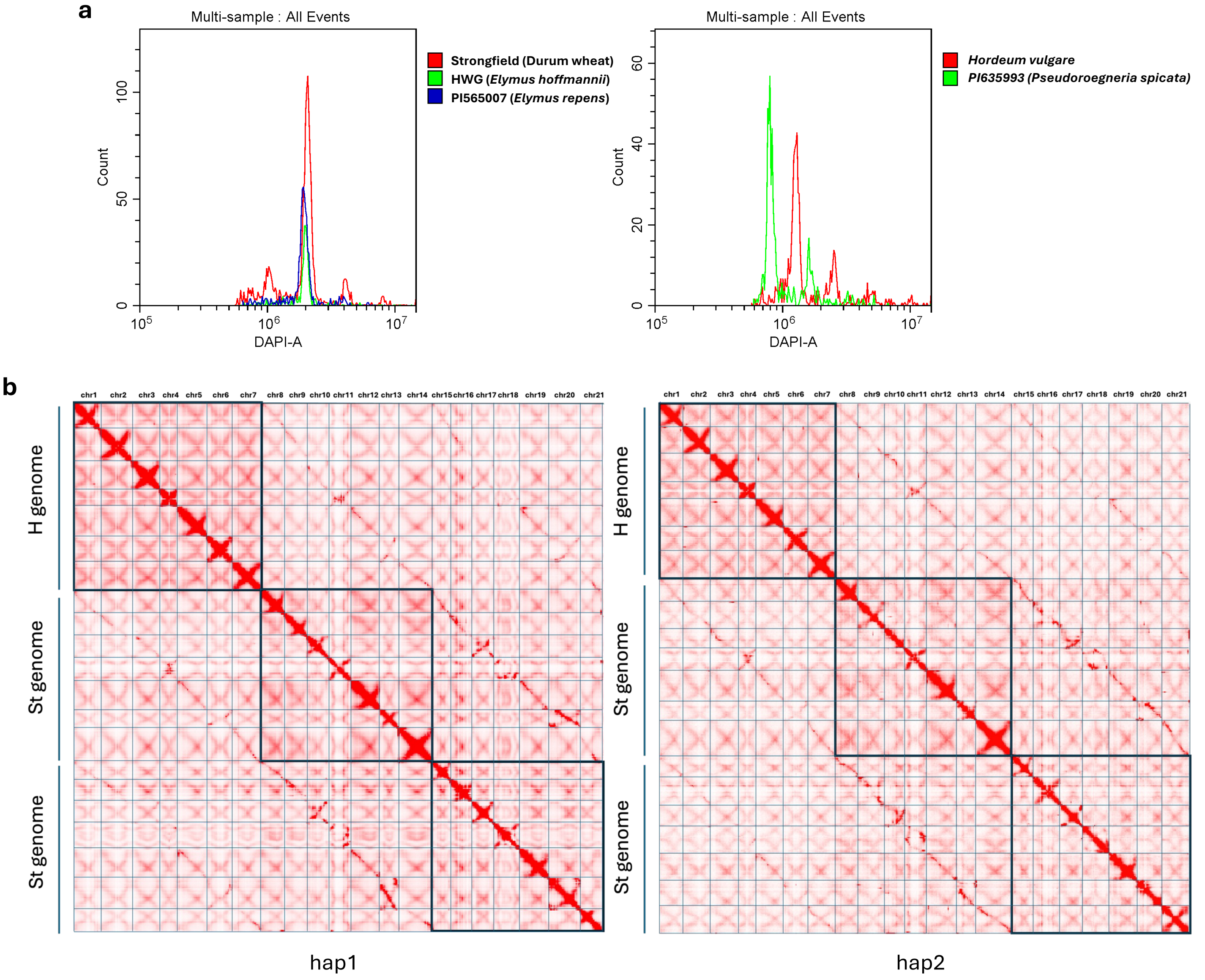


**Supplementary Figure 2 | Genome size estimation and Hi-C contact map. a**. Genome size estimation by flow cytometry. Durum wheat: ~ 12 Gb, HWG: ~11.12 Gb, *Elymus repens:* 10.45 Gb, *P. spicata*: 3.45 Gb. **b.** Hi-C interaction maps of hap1 and hap2 of HWG. Note: The densities of interaction signals were colored in red.

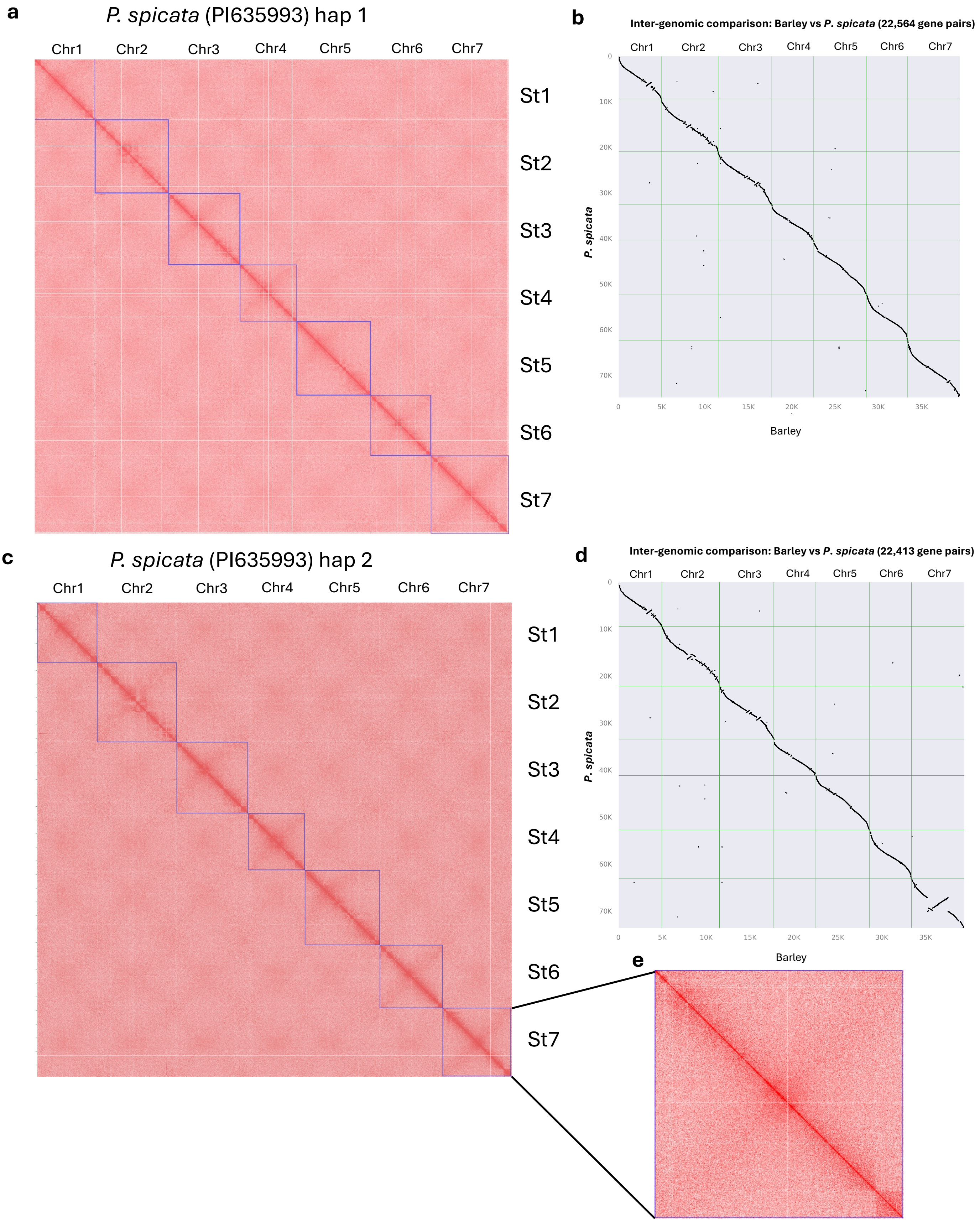


**Supplementary Figure 3 | a and c.** Hi-C interaction maps of hap1 and hap2 of *P. spicata* (PI635993). **b and d**. Collinearity analysis comparing *P. spicata* (hap1/hap2) homologous chromosomes and barley chromosomes. **e**. An enlarged Hi-C contact map of chromosome 7 presents detailed visualization of local chromatin interaction features.

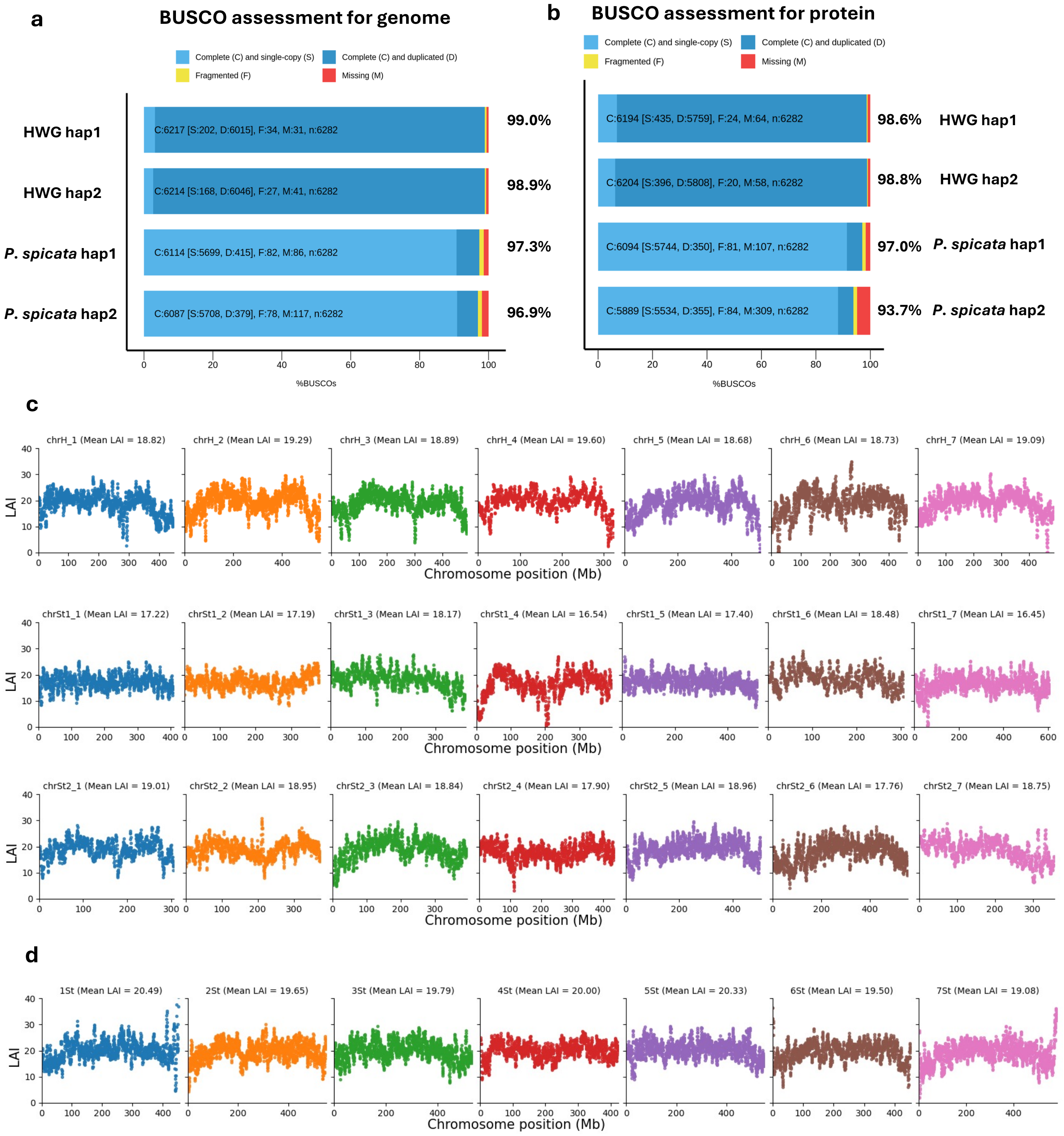


**Supplementary Figure 4 | Genome assembly and annotation quality assessment. a.** BUSCO assessment for assembled genomes. **b**. BUSCO assessment for annotated protein sequences. **c** and **d**. The long terminal repeat (LTR) assembly index (LAI) scores of HWG genome (hap1) (**c**) and *P. spicata* (hap1) (**d**).

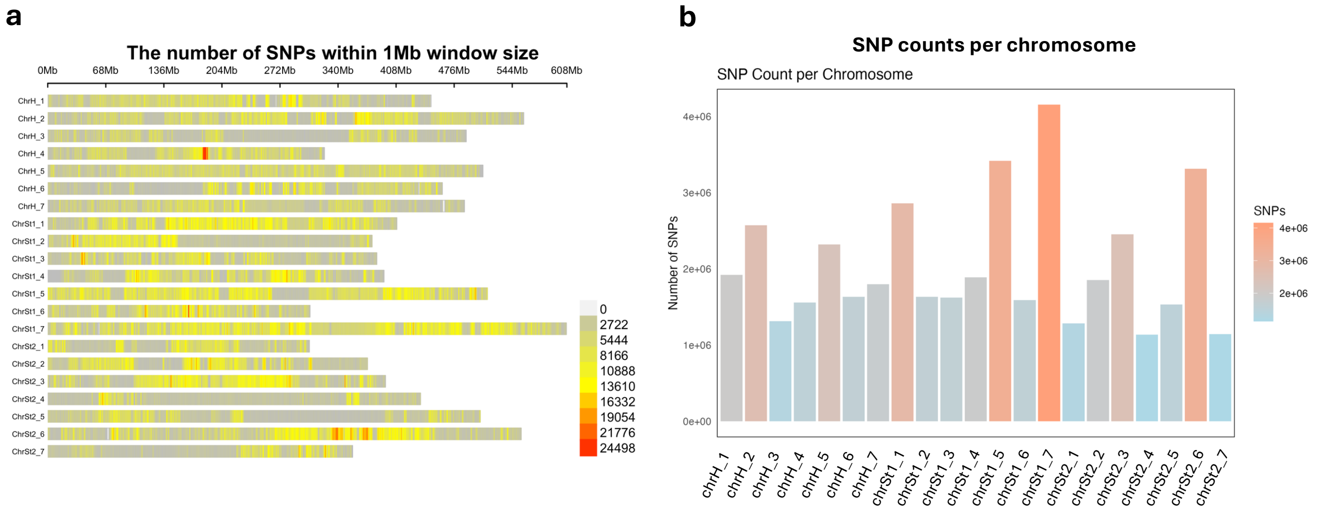


**Supplementary Figure 5 | SNPs distribution indicates heterozygous site in HWG. a.** Density of SNPs within 1 Mb windows size, the density of SNPs were scaled with colors from grey to red. **b**. SNP counts for each chromosomes.

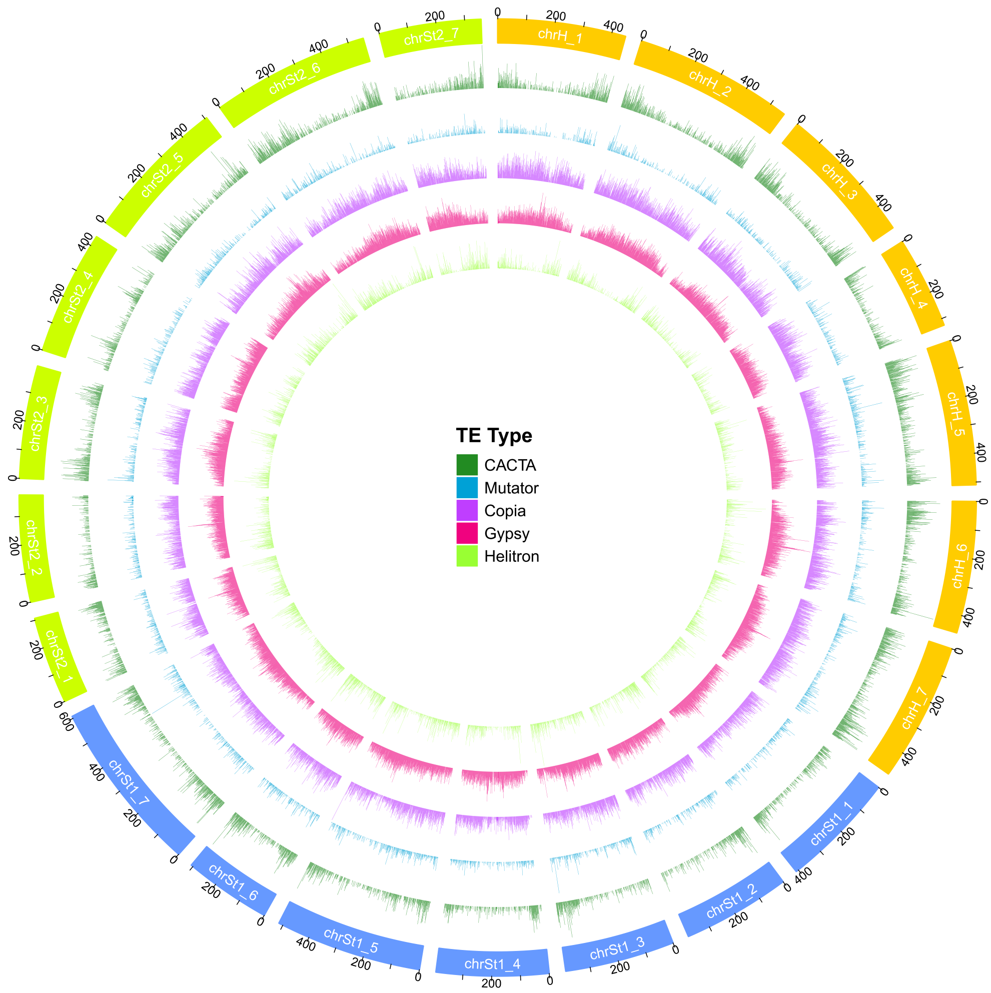


**Supplementary Figure 6 | Overview of TE distribution across the genome of HWG**. Each type of TE were colored in different.

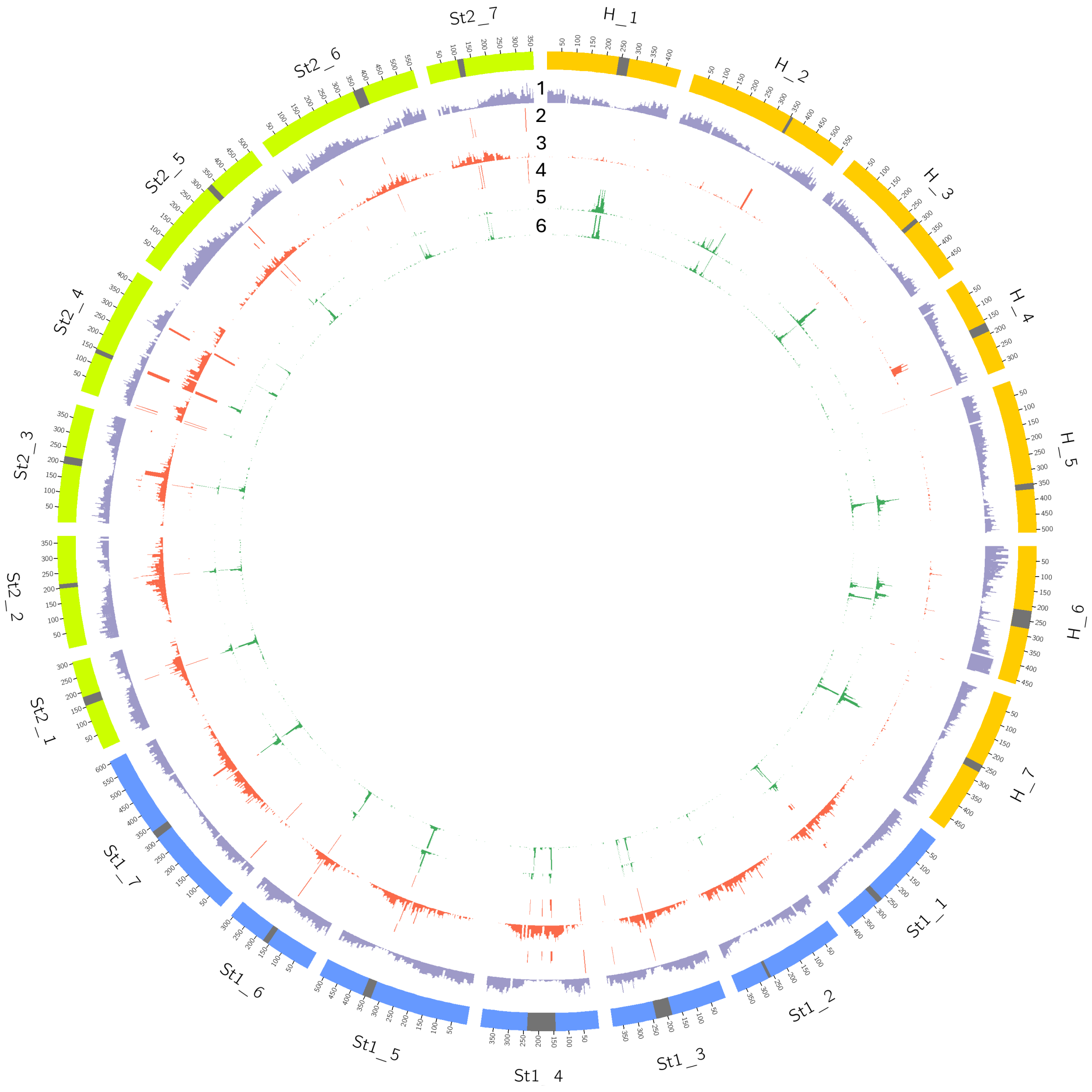


**Supplementary Figure 7 | Centromere region identification**. Track 1: Gene density. Track 2: STlib_96 probe density. Track 3: STlib_98 probe density. Track 4: STlib_117 probe density. Track 5: BAC7 probe (AY040832) density. Track 6: Centromeric retrotransposon of Maize (CRM) TE distribution. Probe STlib_96, STlib_98, and STlib_117 were characterized in the centromeric region in diploid *Pseudoroegneria libanotica* (PI 228392) and were known as St genome specific probes. BAC 7 probe is barley specific probe that located in the centromeric region.

a

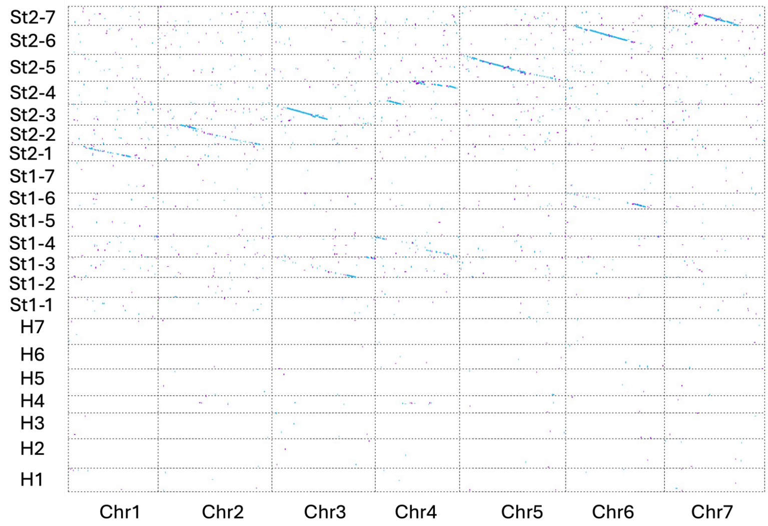


b

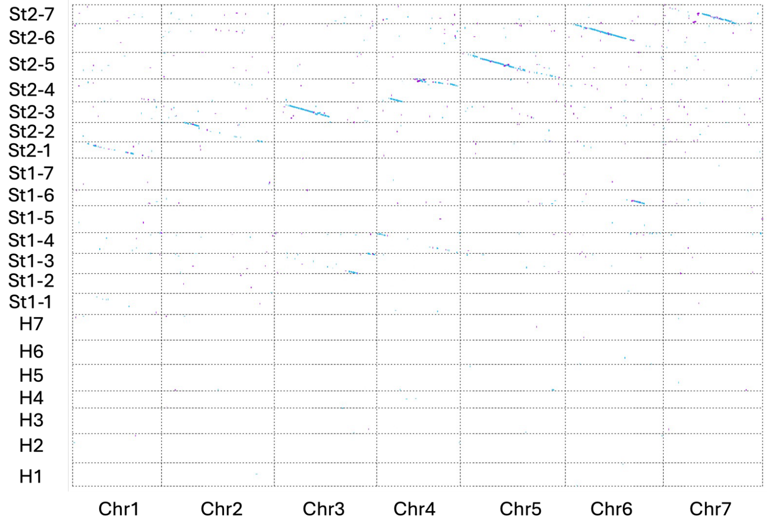


**Supplementary Figure 8 | Subgenome assignments in hexaploid HWG using *P. spicata* as reference.** Dot plots show the maximal exact matches of certain minimum length between two homologous chromosomes. **a**. minimum length is 200 bp. **b**. minimum length is 300 bp.

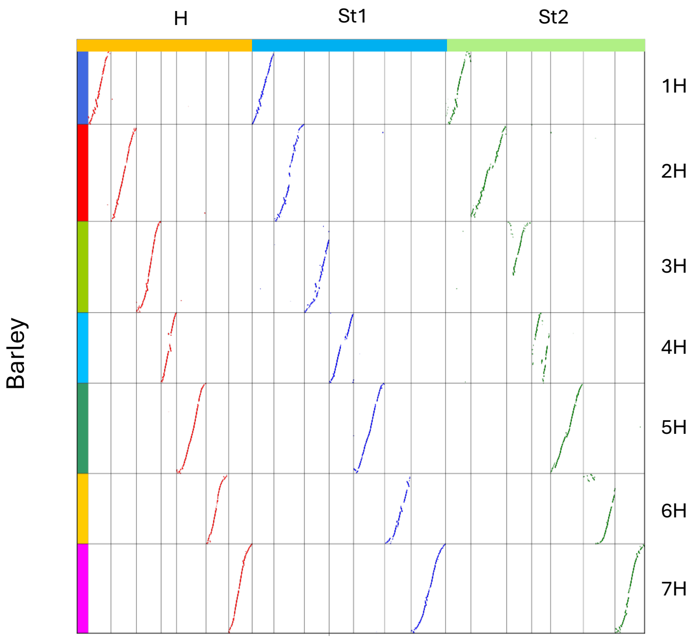


**Supplementary Figure 9 |** Collinearity analysis comparing HWG (hap2) homologous chromosomes and barley chromosome.

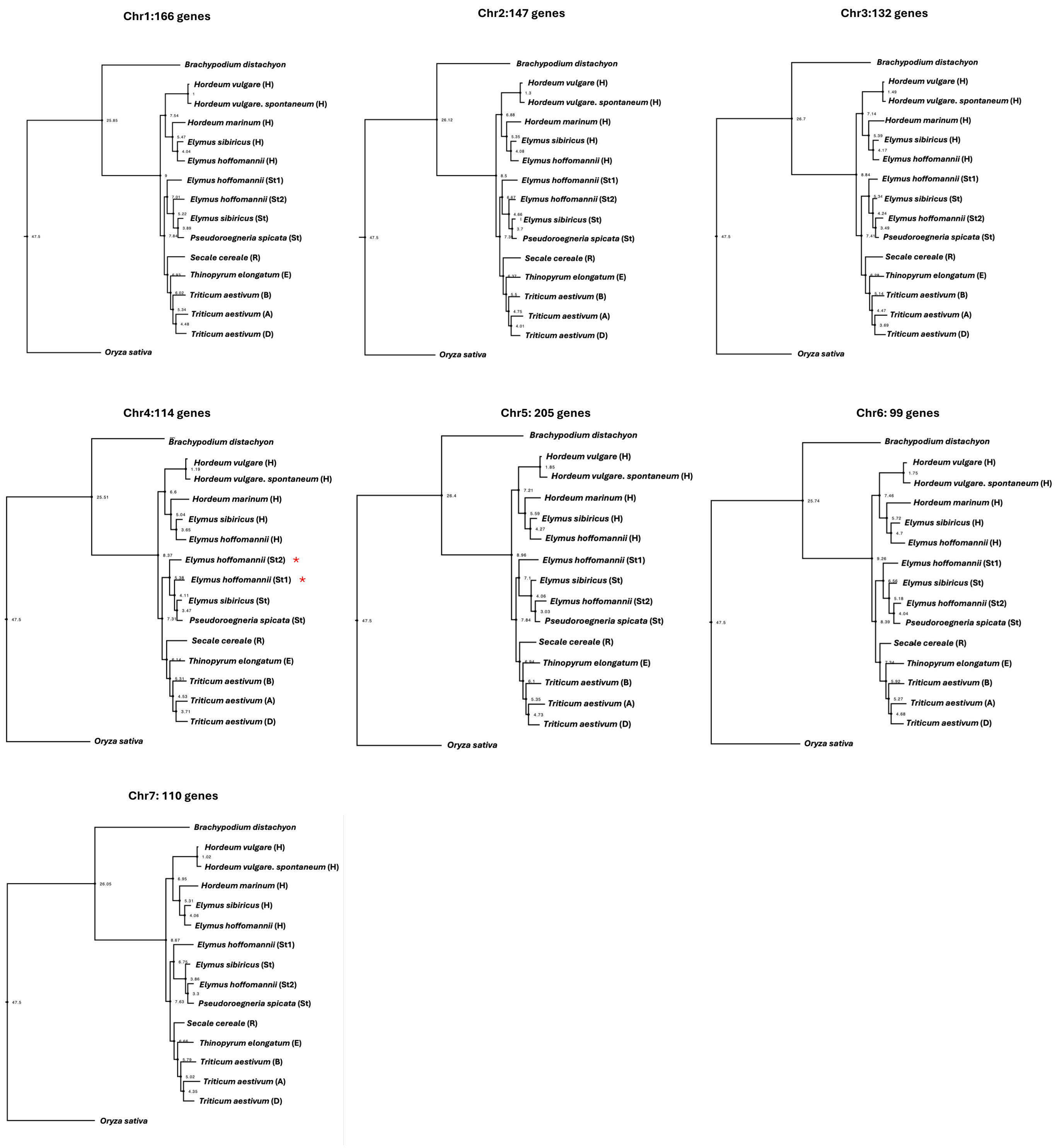


**Supplementary Figure 10 | Assessment of phylogenetic relationship among individual chromosomes based on single-copy genes.** Phylogenetic tree topology inferred from the single-copy genes in the same chromosome using maximum likelihood method. The gene numbers for each chromosome were shown on the top of each tree. Red asterisks indicate the Chr4 showing distinct divergence pattern comparing to other chromosomes.

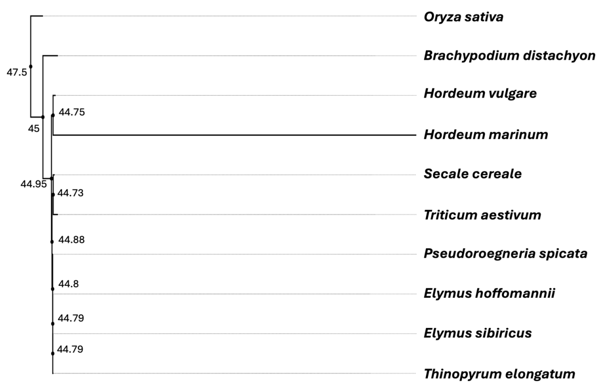


**Supplementary Figure 11 | Phylogenetic relationship among 10 species in Triticeae family based on chloroplast genome sequences.** The divergence time is indicated by each node.

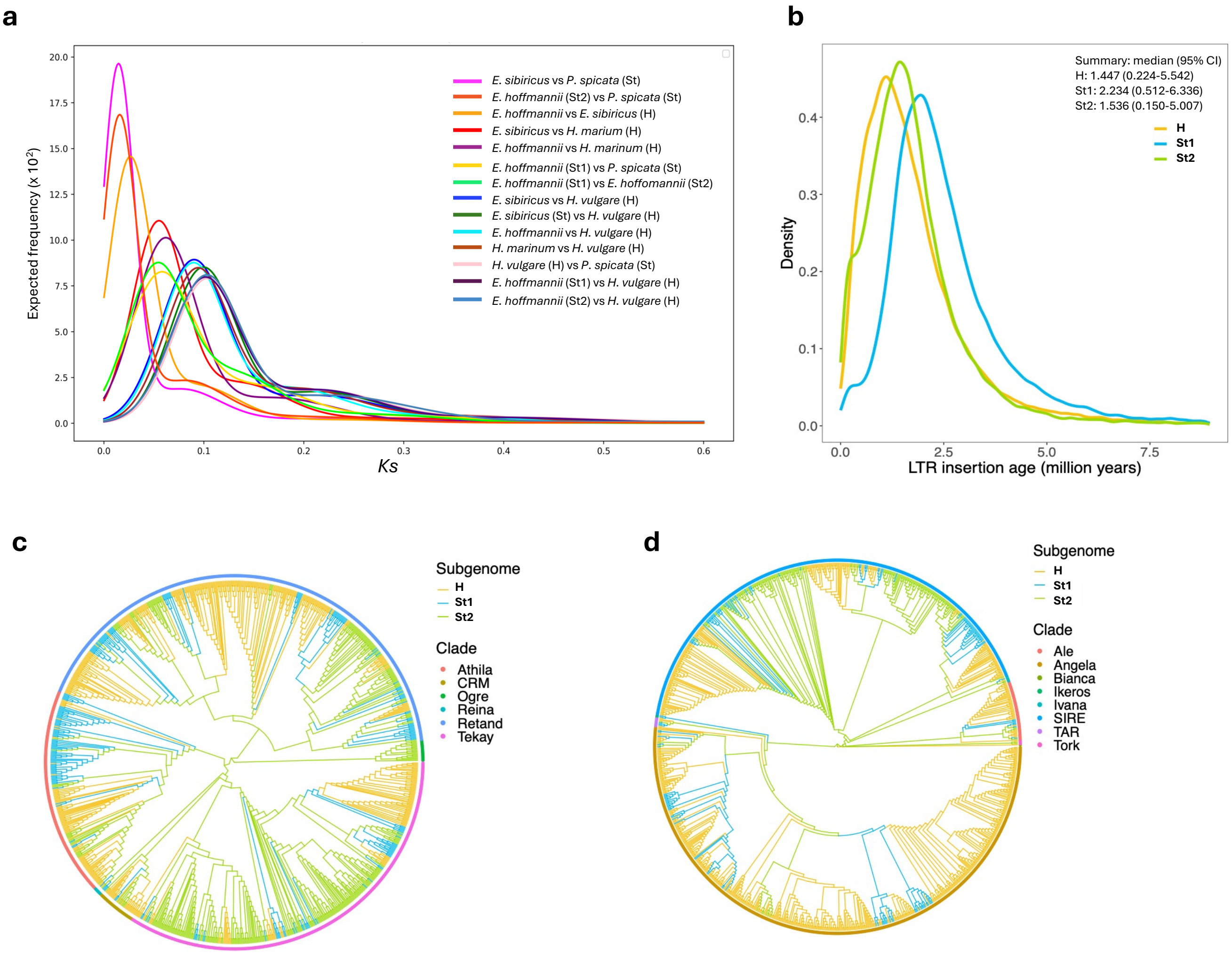


**Supplementary Figure 12 | Divergence time estimation. a.** *K*s frequency distributions. Gaussian mixture models were fitted to the frequency distributions of *Ks* values derived from pairwise comparisons of syntologs between different genomes of each species. **b**. Insertion time of subgenome-specific LTRs. **c**. A phylogenetic tree of 1,000 randomly subsampled LTR/Copia elements. **d**. A phylogenetic tree of 1,000 randomly subsampled LTR/Gypsy elements.

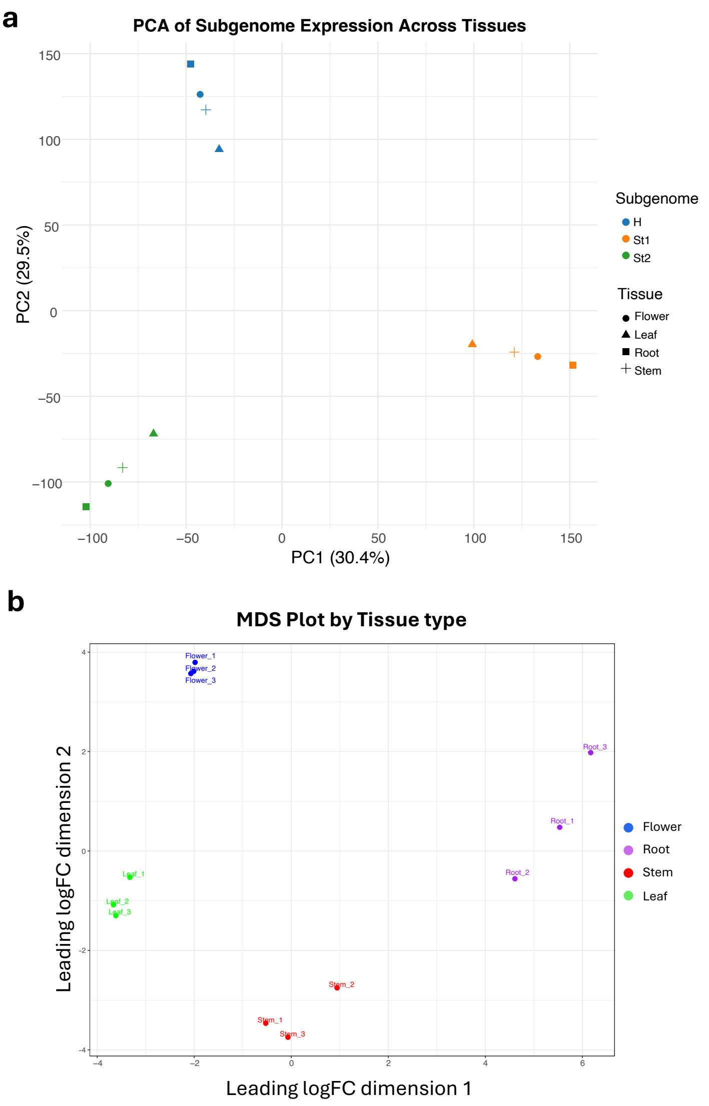


**Supplementary Figure 13 | RNA-seq reads validation.** Principal component analysis (PCA) plots (**a**) and multidimensional scaling (MDS) plots for expression similarity of subgenomes in four tissues (Stem, Flower, Leaf, Root) of HWG.

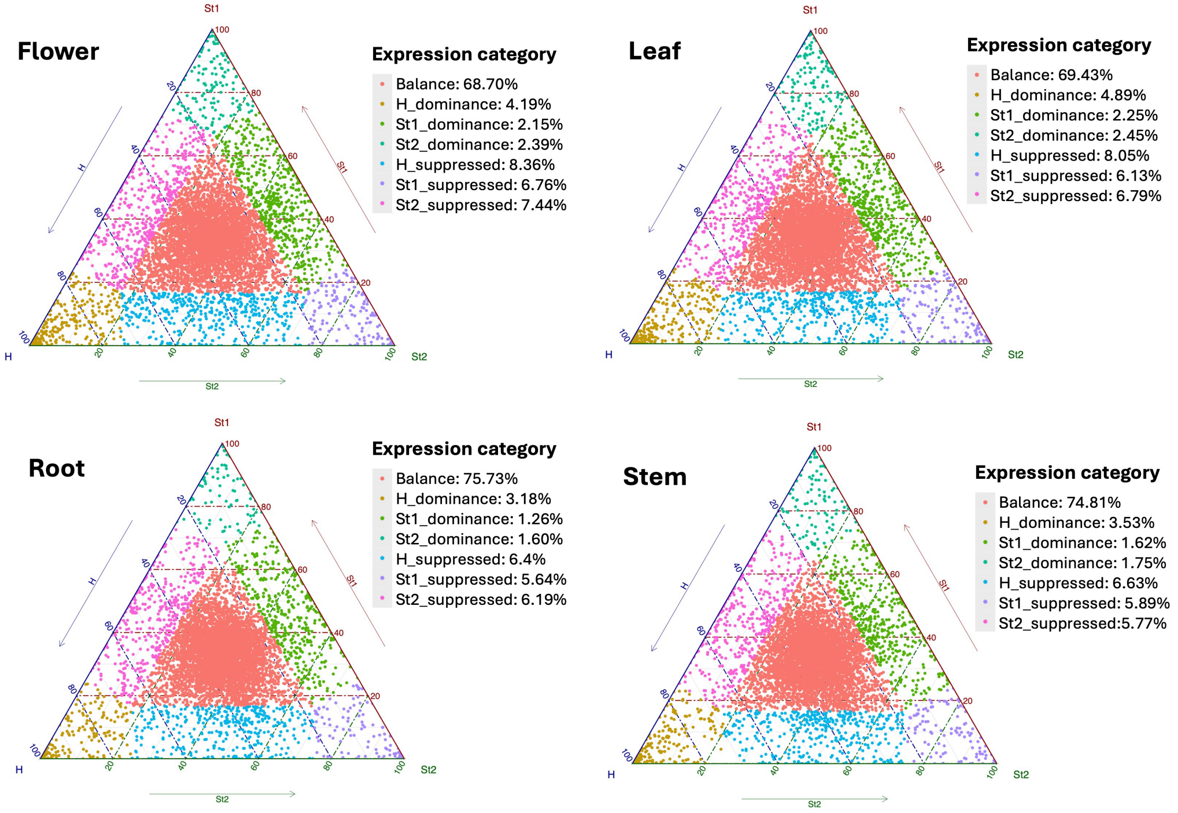


**Supplementary Figure 14 | Homoeolog expression patterns in polyploid HWG. a.** Ternary plot of homoeologs for expression bias across the three subgenome of HWG in different tissues. Each point in the ternary plot represents a gene triad with coordinates corresponding to the H, $St_1$ and $St_2$ subgenomes. Triads in vertices indicate dominant categories, whereas triads near edges or between vertices are suppressed categories; balanced triads shown in the center in red.

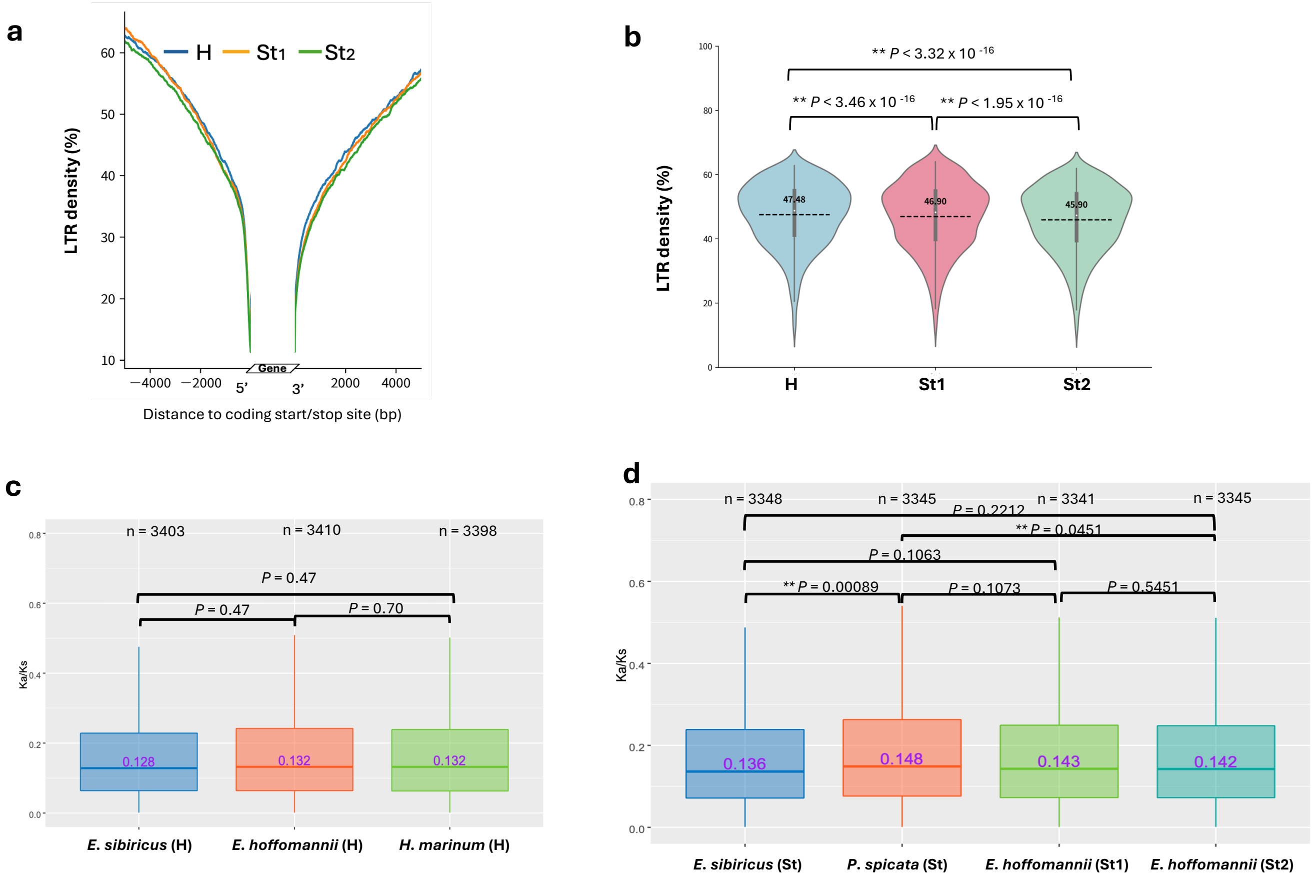


**Supplementary Figure 15 | LTR density and *Ka/Ks* ratios across the subgenomes in HWG. a.** Variation of LTR density in the flanking regions among three subgenomes of HWG. **b.** LTR density distribution patterns among subgenomes of HWG. The asterisks represent significant differences (based on two-tailed Wilcoxon rank-sum test). **C and d.** Box plots comparing *Ka/Ks* value distributions among the H subgenomes and St subgenomes in *E. sibiricus*, HWG and *H. marinum*. The central line for each box plot denotes the median. Sample size (n) used in each comparison are indicated. The asterisks denote significant differences based on two-tailed Wilcoxon rank-sum test.

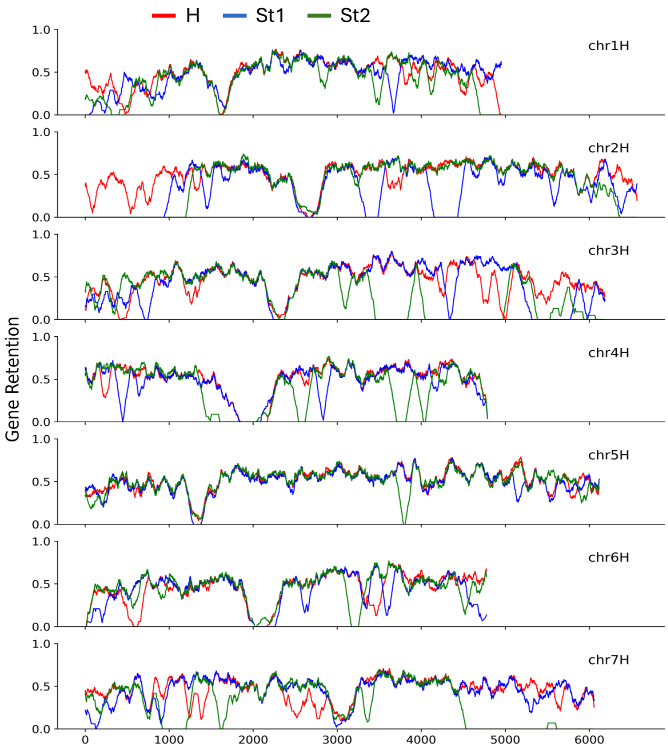


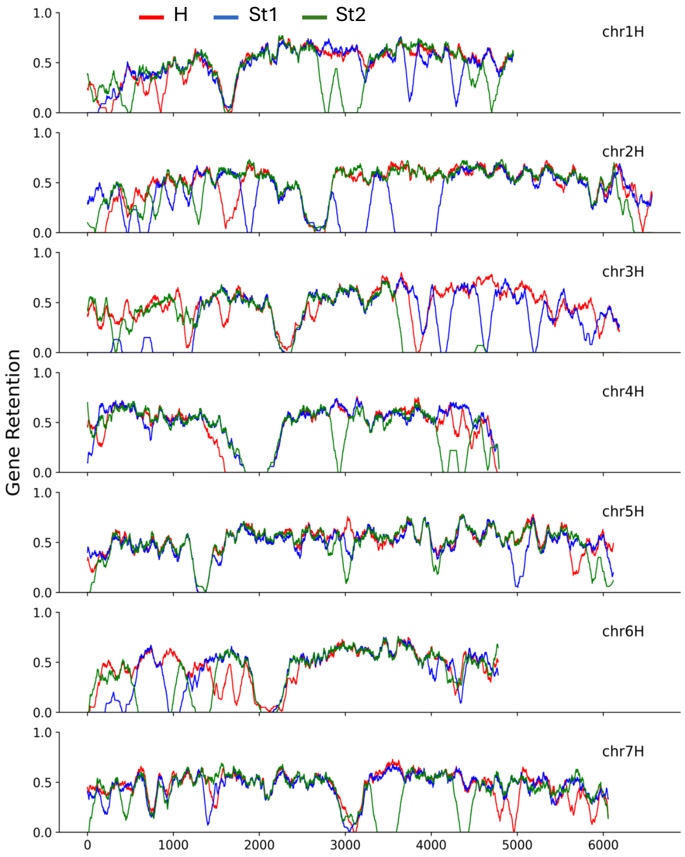


**Supplementary Figure 16 | Gene retention analysis in hap1 (upper) and hap2 (bottom) of HWG**. Gene retention patterns among subgenomes of HWG relative to the barley genome.

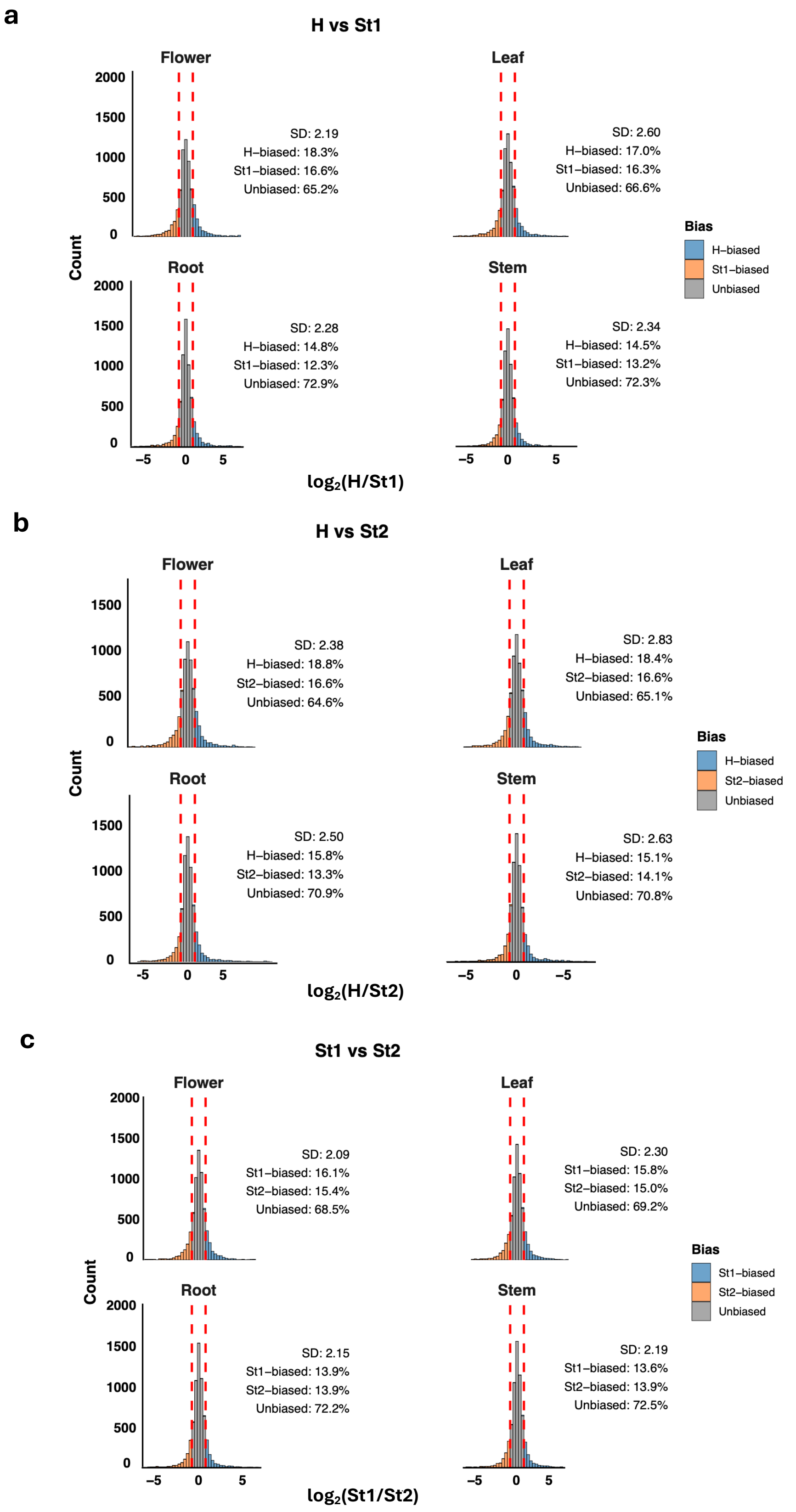


**Supplementary Figure 17 | Homoeolog expression patterns in different tissues of polyploid HWG.** Histograms showing the number of up-regulated genes across four tissues in HWG in each subgenome comparisons (a. H vs $St_1$, b. H vs $St_2$, c. $St_1$ vs $St_2$).

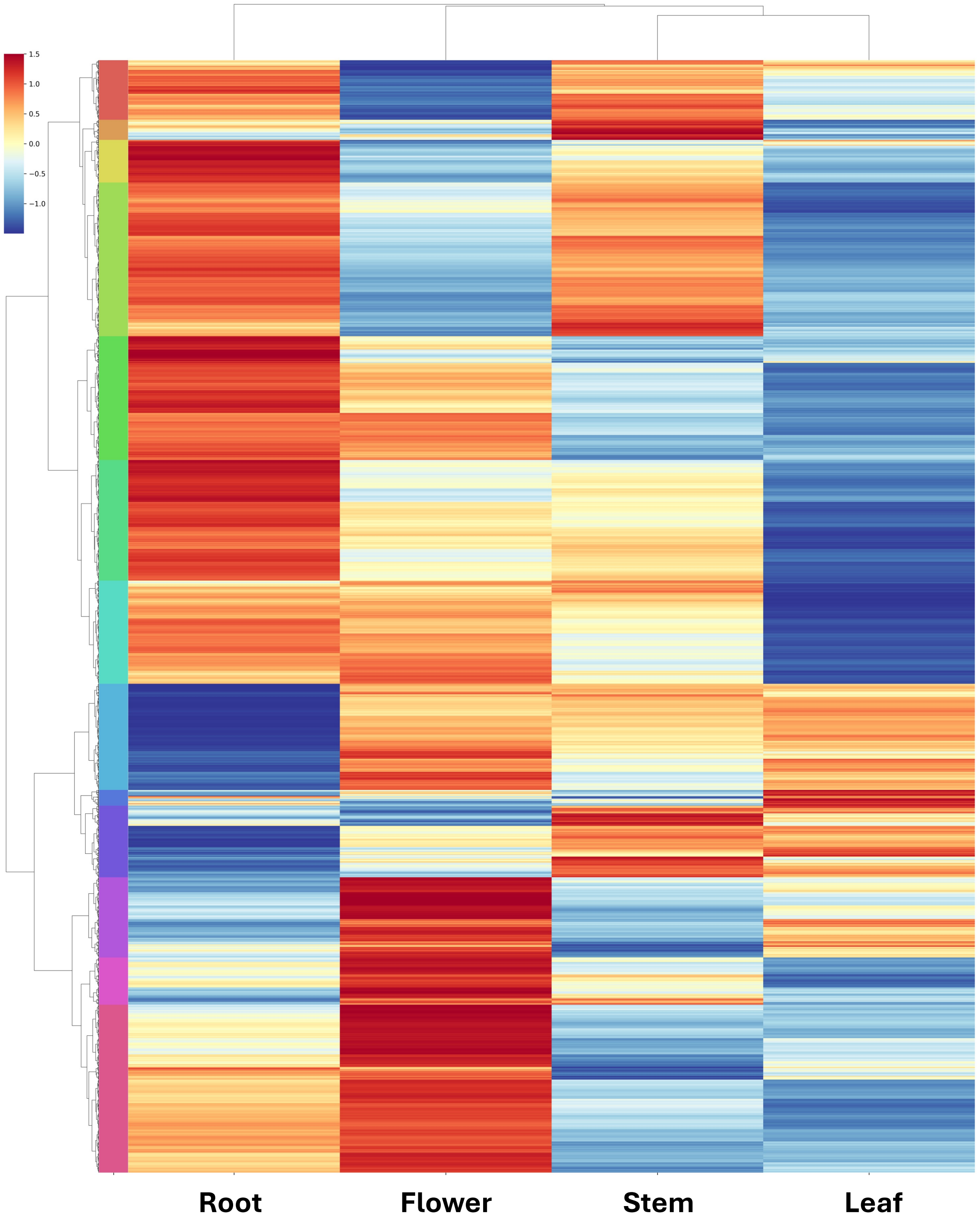


**Supplementary Figure 18 | The correlation of gene expression between homoeologs across four tissues in HWG.** 6742 genes were clustered into 13 groups indicated by different color bars on the left.
